# A di-arginine additive for dissociation of gold nanoparticle aggregates: A matrix-insensitive approach with applications in protease detection

**DOI:** 10.1101/2022.09.30.508454

**Authors:** Maurice Retout, Zhicheng Jin, Jason Tsujimoto, Yash Mantri, Raina Borum, Matthew N. Creyer, Wonjun Yim, Tengyu He, Yu-Ci Chang, Jesse V. Jokerst

## Abstract

We report the reversible aggregation of gold nanoparticle (AuNPs) assemblies via a diarginine peptide additive and thiolated PEGs (HS-PEGs). The AuNPs were first aggregated by attractive forces between the citrate-capped surface and the arginine side chains. We found that HS-PEG thiol group has higher affinity for the AuNPs surface, thus leading to redispersion and colloidal stability. In turn, there was a robust and obvious color change due to on/off plasmonic coupling. The assemblies’ dissociation was directly related to the HS-PEG structural properties such as their size or charge. As an example, HS-PEGs with a molecular weight below 1 kDa could dissociate 100% of the assemblies and restore the exact optical properties of the initial AuNPs suspension (prior to the assembly). Surprisingly, the dissociation capacity of HS-PEGs was not affected by the composition of the operating medium and could be performed in complex matrices such as plasma, saliva, bile, urine, cell lysates or even sea water. The high affinity of thiols for the gold surface encompasses by far the one of endogenous molecules and is thus favorized. Moreover, starting with AuNPs already aggregated ensured the absence of background signal as the dissociation of the assemblies was far from spontaneous. Remarkably, it was possible to dry the AuNPs assemblies and to solubilize them back with HS-PEGs, improving the colorimetric signal generation. We used this system for protease sensing in biological fluid. Trypsin was chosen as model enzyme and highly positively charged peptides were conjugated to HS-PEG molecules as cleavage substrate. The increase of positive charge of the HS-PEG-peptide conjugate quenched the dissociation capacity of the HS-PEG molecules which could only be restored by the proteolytic cleavage. Picomolar limit of detection was obtained as well as the detection in saliva or urine.

**TOC:** 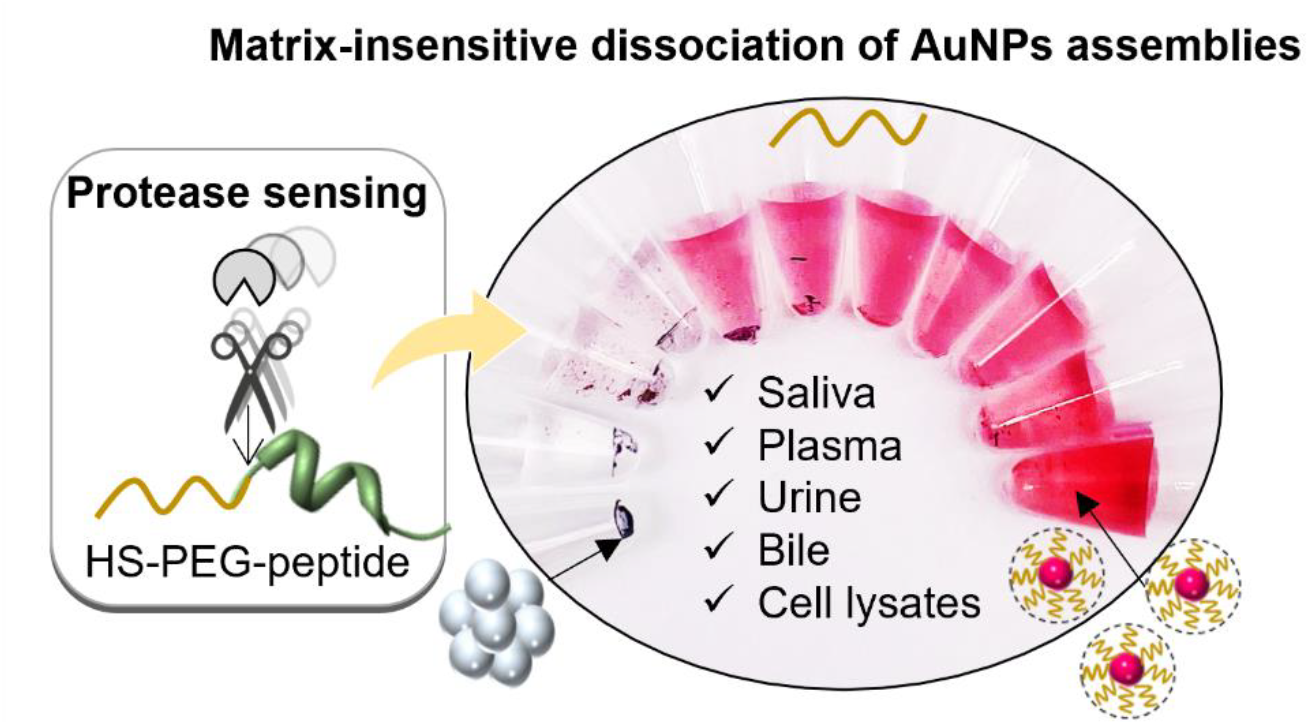

## INTRODUCTION

Noble nanomaterials and particularly gold nanoparticles (AuNPs) have emerged as remarkable colorimetric reporters due to their plasmonic properties.^1–3^ AuNPs possess a localized surface plasmon resonance (LSPR) band in the visible region that is modulated by their size, shape and dielectric environment.^4–6^ Citrate-capped AuNPs (AuNPs-citrate) are the most common, convenient, and cost-effective form of gold colloids, and their synthesis *via* the Turkevich method is well established in the literature.^7^ AuNPs have been extensively studied, and various AuNPs-based colorimetric sensors have been developed.^8–10^

One attractive and common sensing/analysis strategy with AuNPs is the cross-linking of the particles induced by the analyte that couple individual LSPR bands, thus shifting the initial red color of the dispersed AuNPs to blue as the distance between particles diminishes.^8,11,12^ Different mechanisms have been thus exploited to induce the assembly of AuNPs such as DNA pairing,^13^ antibody-antigen interactions,^10^ electrostatic interactions^8^, and hydrogen bonding.^14^

However, one limitation is their colloidal stability. Indeed, their dispersibility is inherent to the balance of attractive Van der Waals (VdW) and repulsive electrostatic forces.^15^ Thermodynamically, AuNPs colloids are not stable and tend to aggregate over time, which results in non-desired aggregation of the particles that reduces the color difference between a positive sample (analyte present) and a negative sample (analyte absent).^8,16^ Another limitation is the difficulty to operate in biofluids such as plasma, saliva, urine, bile or cell lysates. The presence of background matrix compounds can interfere with the colorimetric detection, e.g., via a protein corona that prevents the AuNPs cross-linking.^17^ Also, dramatic background signals are usually observed in biofluids because there is significant charge screening, thus leading to a reduction in repulsive forces and aggregation of the AuNPs due to the VdW interaction. Extreme dilution of the biological sample ^16^ or extraction of the analyte from the matrix are thus often needed.^18^ Only a few examples of nanoparticle aggregation-based assays in biological fluids have been reported.^19,20^

Here, we show a novel process of reversible aggregation of AuNPs-citrate for alternative sensing strategies. Reversible aggregation of nanoparticles is challenging because, according to the Derjaguin-Landau-Verwey-Overbeek (DLVO) theory,^21^ the particles can be trapped in a deep energetic minima during the aggregation, thus transforming the aggregates into larger insoluble materials that can be only slightly dissociated by aggressive sonication; their optical properties cannot be recovered.^22^ While many in the community have shown reversible assembly of a small number of nanoparticles, this usually requires sophisticated coating of the AuNPs-citrate surface with responsive polymers, ^23,24^ DNA strands ^25,26^ or other organic ligands.^27,28^ However, we are unaware of work describing the assembly of large amount of AuNPs-citrate into macroscopic aggregates that can easily be dissociated without the need for prior surface modifications.

Our system uses only a di-arginine additive that causes the aggregation of AuNPs and thiolated poly(ethylene glycol) (HS-PEGs) for the dissociation of the assemblies (**Scheme 1**). The system works through differences in affinity of the surface ligands: the introduction of HS-PEGs leads to redispersion of the AuNPs due to the higher affinity of the thiol for the gold surface than the arginine or citrate. In turn, there is a plasmonic color change. Starting with assembled AuNPs allows for insight on the colorimetric signal. Surprisingly, the redispersion can be performed over a wide variety of solvents including human plasma, serum, saliva, and even bile. After characterizing the ligands and HS-PEGs best suited for this reaction, we deployed it for colorimetric sensing of proteases via a cleave PEG-peptide conjugate. Although the grafting of HS-PEGs molecules on dispersed AuNPs has been widely reported in the literature, to the best of our knowledge, the dissociation of AuNPs assemblies with HS-PEGs has never been studied before, especially for the detection of proteases.

**Scheme 1.**
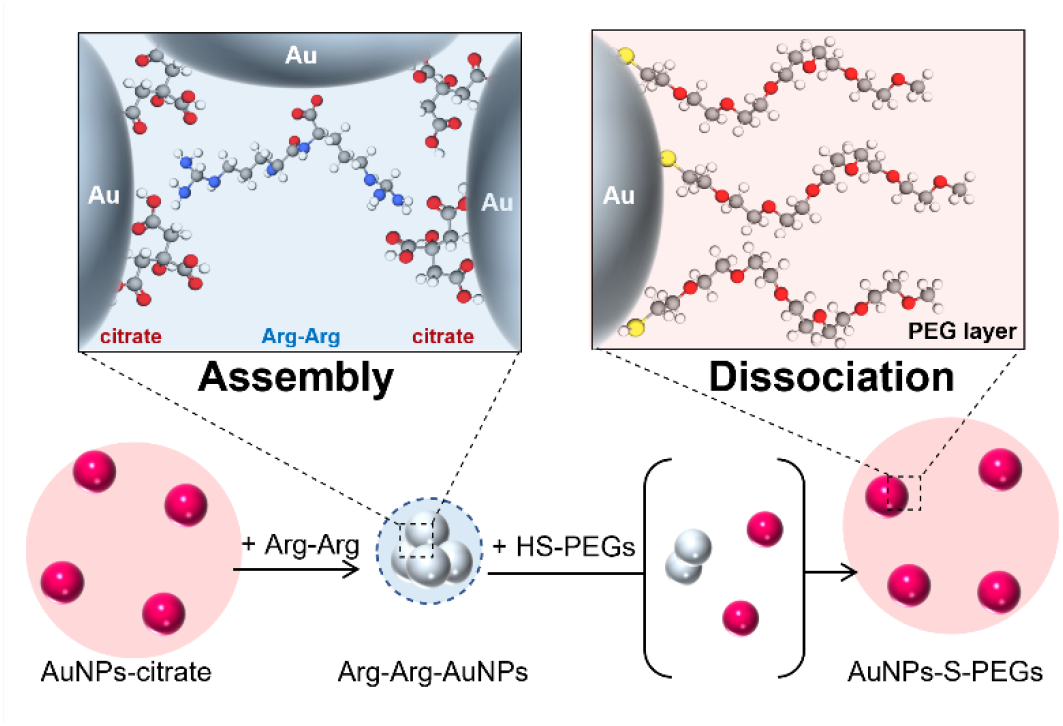
Assembly of citrate-capped AuNPs (AuNPs-citrate) with a di-arginine peptide and subsequent dissociation with thiolated PEG molecules (HS-PEGs).

## RESULTS AND DISCUSSION

### Formation of AuNP assemblies

In this study, we investigated the possibility of forming convenient and reversible AuNPs assemblies using only elementary 15 nm AuNPs-citrate suspended in water and short peptides without the need for a complex surface modification. Peptides were used because they are relatively bulky and thus their steric prevented the particles from entering a permanent aggregated state. Arginine in particular can strongly interact with citrate anions via electrostatic interactions. A dipeptide containing two repetitions of arginine (Arg-Arg or RR) was thus used to interact with multiple particles at the same time and induce assembly.

Transmission electronic microscopy (TEM) revealed that the addition of excess (10^4^ equiv.) of di-arginine peptide (Arg-Arg) to an aqueous suspension of AuNPs-citrate led to bulky assemblies and no dispersed AuNPs were observed (**Figure 1A** vs **1B**). Multispectral advanced nanoparticles tracking analysis (MANTA) was then used to confirm the size increase: The initial blue scattering corresponding to a hydrodynamic diameter of 40 ± 25 nm (**Figure 1C**) transformed immediately into red scattering corresponding to a hydrodynamic diameter of approximately 467 ± 120 nm (**Figure 1D**). Importantly, the count of particles decreased by more than 90%—from 2.5×10^5^/mL to 2×10^4^/mL, thus confirming that the vast majority of the AuNPs-citrate were aggregated. TEM images at different magnifications are seen in the supporting information (Figure S1 and S2).

**Figure 1.**
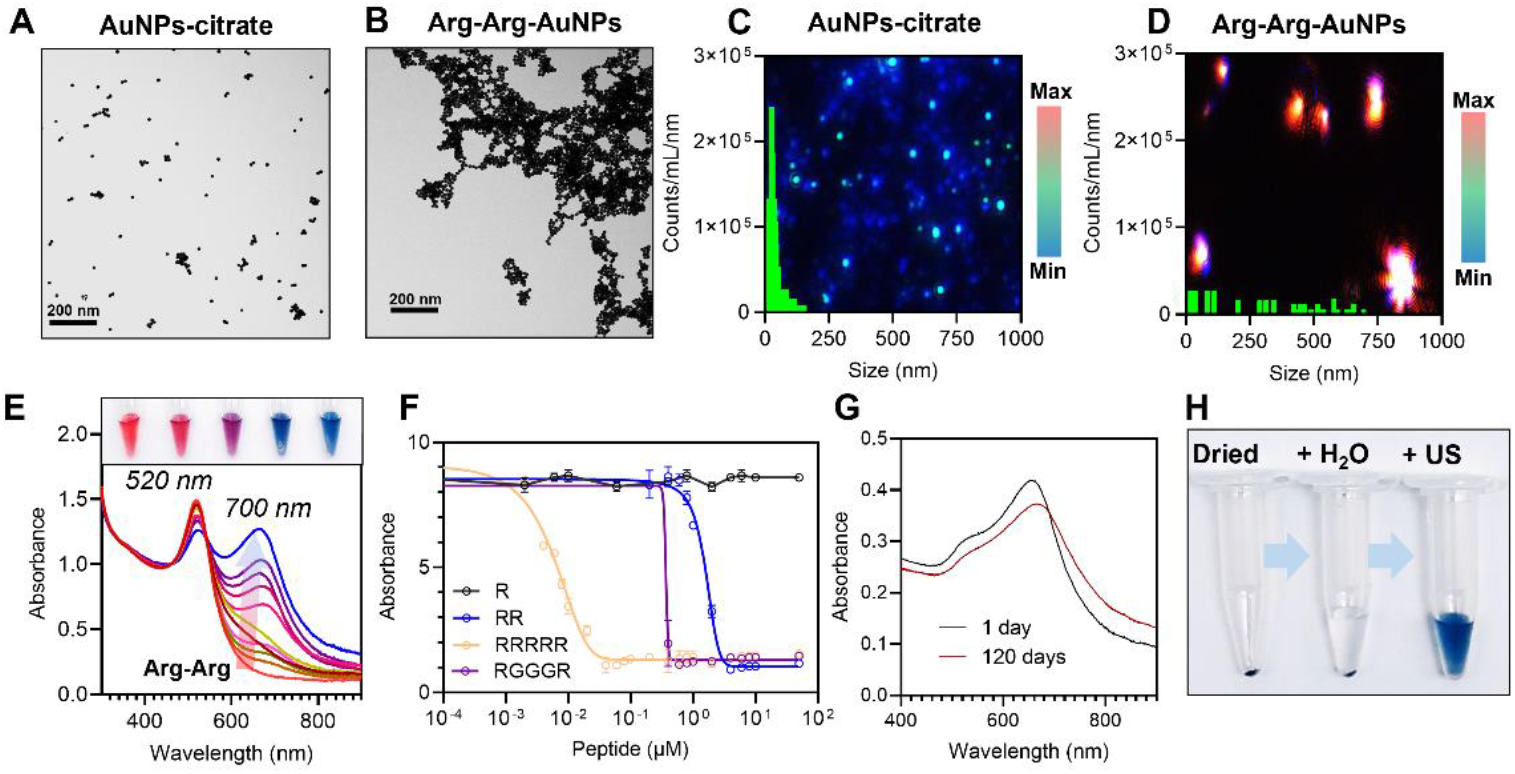
Peptide-induced assembly of AuNPs-citrate. **(A, B, C and D)** TEM images and multispectral advanced nanoparticle tracking analysis (MANTA)^29^ image and size distribution of AuNPs-citrate before **(A and C)** or 10 minutes after the addition of 10 μM of Arg-Arg **(B and D)**. MANTA records images of the particles’ light scattering via three differently colored lasers (e.g., blue, green, and red); the scattering depends on the particle’s size. MANTA counts the nanoparticles and calculates their size. **(E)** Modification of the optical properties of AuNPs-citrate during the assembly with Arg-Arg. Insets show the pictures of the samples for concentrations of Arg-Arg of 0, 1, 2, 5 and 10 μM. **(F)** Ratio of the absorbances Abs.520nm/Abs.700nm of AuNPs-citrate as a function of peptides concentration. **(G)** UV-Vis spectra of Arg-Arg-AuNPs 1 day or 120 days old. **(H)** Pictures of Arg-Arg-AuNPs dried and resuspended in pure water after 5 seconds of sonication.

The peptide-induced assembly of AuNPs-citrate strongly impacted the optical properties of the colloid. An immediate modification of the LSPR band of the particles was observed proportional to the peptide concentration. The absorbance at λ_max_ decreased, and a new absorption peak increased at 700 nm (**Figure 1E**). Such deformation of the LSPR band led to a change of color of the suspensions turning from bright red to blue (**Figures 1E**, insets). The ratio of the absorbance at 520 nm over 700 nm were then used to characterized the AuNPs assemblies through all this study. Interestingly, the density of Arg-Arg leading to the maximal ratio of absorbance was approximately 1.8 Arg-Arg-/nm^2^, which corresponds approximately to the maximal density of Arg-Arg that could be carried by one AuNPs (1.6 Arg-Arg/nm^2^) considering a footprint for the peptide of 0.63 nm^2^ (see http://biotools.nubic.northwestern.edu/proteincalc.html).

Arg-Arg was chosen to promote the AuNPs-citrate assemblies because it is the minimal sequence that can induce the assembly. The reversibility of our system is based on the replacement of citrate and peptide layers by PEG, and it was thus crucial to remove the peptides from the AuNPs surface. While a single arginine could not induce any form of assembly, peptides containing more than two arginine had a higher affinity for the AuNPs-citrate than Arg-Arg and were thus discarded (**Figure 1F**). As a control, we synthesized a peptide containing two arginine but spaced by three glycine (RGGGR) to decrease the charge density and thus lower the affinity of the peptide for the AuNPs-citrate. However, RGGGR had a higher affinity for the particles than RR (Arg-Arg) and was also discarded.

Despite its short length, Arg-Arg was sufficient to protect the AuNPs from degradation, and the resulting assemblies were highly stable over time. **Figure 1G** shows barely no difference between the LSPR band of pristine and 120-day-old Arg-Arg-AuNPs. To our delight, another demonstration of the high stability of the assemblies was performed with their drying and resuspension in pure water using only 5 seconds of sonication without any degradation or loss of particles (**Figure 1H**).

To demonstrate the versatility of our system, this study was reproduced with 40 nm AuNPs-citrate commercially available from Nanocomposix (supporting information Section 7).

### Dissociation of the AuNPs assemblies

The proof-of-concept of the assembly reversibility was demonstrated with a polyethylene glycol (PEG) system containing six repetitions capped with a methoxy group and a thiol at the other end (HS-PEG_6_-OCH_3_). HS-PEG_6_-OCH_3_ was chosen because its grafting on AuNPs ensure a high colloidal stability of these latter. Indeed, the grafting of thiolated PEGs has been widely reported in the literature via the formation of a strong Au-S bond conferring to the particles a high colloidal stability due to a combination of hydrophilicity and steric hindrance.^30–32^ The Au-S bond being stronger than the arginine-citrate, arginine-gold or even citrate-gold interactions, HS-PEG_6_-CH_3_ was expected to be capable of progressively replacing the arginine/citrate layers at the gold surface, disrupting the electrostatic network, and eventually dispersing the AuNPs (**Scheme 1**). Briefly, aqueous suspensions Arg-Arg-AuNPs assemblies were exposed to increasing concentrations of HS-PEG_6_-OCH_3_, and the dissociation of the assemblies was characterized by TEM, MANTA, FTIR and UV-Vis spectroscopies (**Figure 2**).

**Figure 2.**
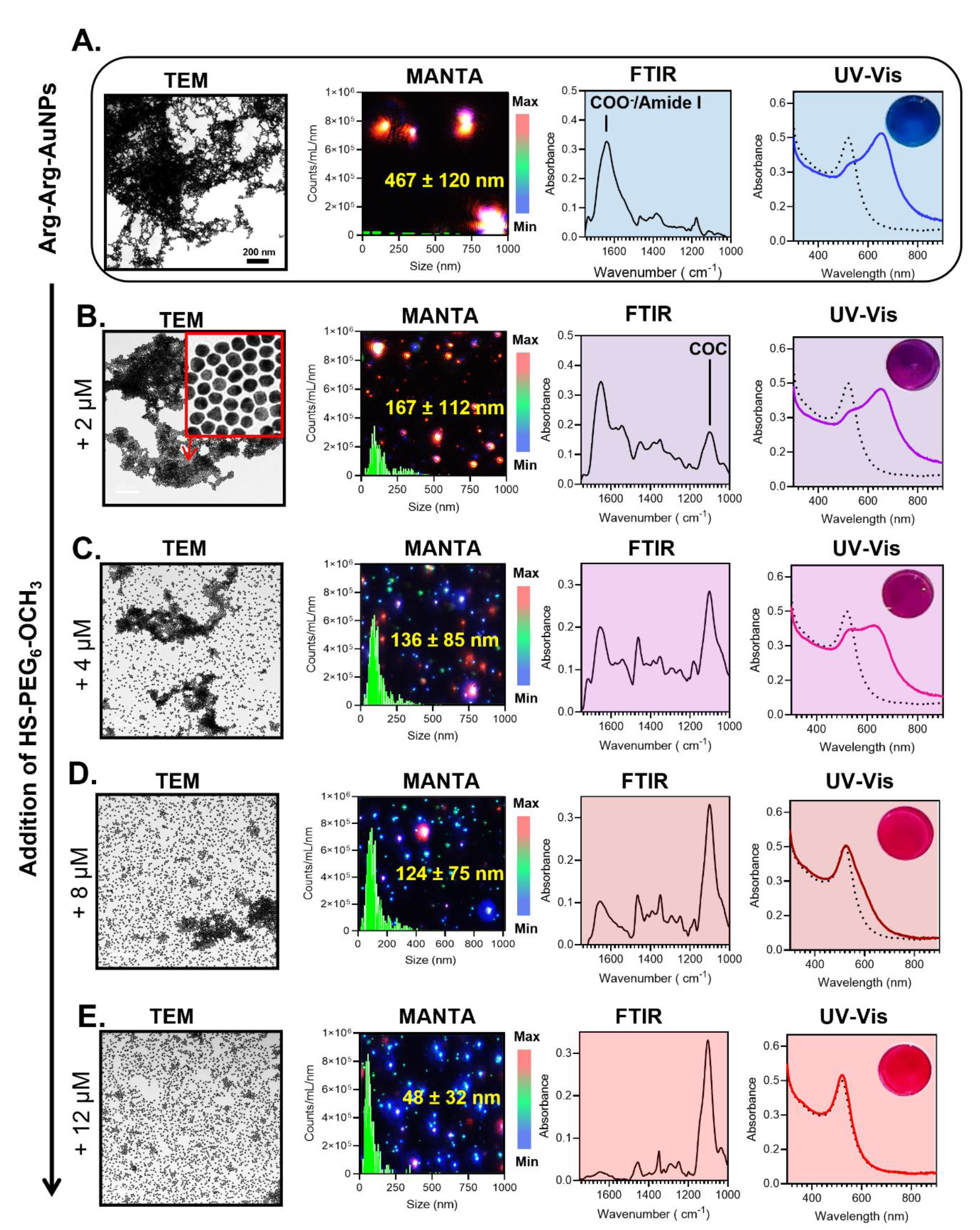
Characterization of the dissociation of AuNPs assemblies with HS-PEG_6_-OCH_3_. From left to right, TEM*, MANTA, ATR-FTIR**, and UV-Vis*** spectroscopy analysis of Arg-Arg-AuNPs **(A)** before or 10 minutes after the addition of **(B)** 2 μM, **(C)** 4 μM, **(D)** 8 μM, and **(E)** 12 μM of HS-PEG_6_-OCH_3_. *All the images are at the same magnification. **All samples were cleaned by centrifugation from non-bound molecules (HS-PEGs, citrate or Arg-Arg) and the FTIR spectrum corresponds only to molecules attached to the gold surface. *** Black dashed line shows the LSPR band of the initial AuNPs-citrate.

Prior to any addition of HS-PEGs, the dense and large aggregates of Arg-Arg-AuNPs present only ATR-FTIR signals coming from citrate and/or Arg-Arg as well as COO^−^_νas_ and/or amide band I around 1650 cm^−1^, respectively. The color of the sample was bright blue because the LSPR band was strongly red-shifted (**Figure 2A)**.

After the addition of 2 μM of HS-PEG_6_-OCH_3_, however, the distance between the AuNPs in the aggregates increased and the size of the aggregates decreased from 467 ± 120 nm to 167 ± 112 nm. Interestingly, ATR-FTIR spectroscopy showed an increase in the absorbance at 1100 cm^−1^ corresponding to the C-O-C stretching of the PEG chain and a decrease in the absorbance around 1650 cm^−1^. This indicates that few HS-PEG_6_-OCH_3_ were grafted onto the AuNPs surface, which explains the increase in the interparticle distance. The decrease in the size of the assembly led to a color change from blue to purple (**Figure 2B** and S3).

**Figures 2C** and **2D** show the results of the addition of 4 μM and 8 μM of HS-PEG_6_-OCH_3_, respectively. More AuNPs detached from the aggregates and became monodisperse with higher concentrations of HS-PEG_6_-OCH_3_ (Figure S4 and S5). Accordingly, the average size of the aggregates decreased to 136 ± 85 nm and 124 ± 75 nm for 4 μM and 8 μM, respectively. Moreover, the intensity of the ATR-FTIR signals of the PEG chain increased and the signal of the citrate/Arg-Arg decreased, thus confirming the expansion of the grafting density of HS-PEG_6_-OCH_3_. The LSPR band of the colloid red-shifted due to dissociation: The solution color were purple-pink and wine red at 4 μM and 8 μM, respectively.

At 12 μM HS-PEG_6_-OCH_3_ and above, aggregates were no longer seen in TEM, and MANTA measured a mean size of 48 ± 25 nm. ATR-FTIR showed only absorbance of the C-O-C signal with limited citrate/Arg-Arg signals, thus indicating that the citrate/arginine layer was completely removed from the surface for the benefit of HS-PEG_6_-OCH_3_. The presence of a PEG layer around the AuNPs explains the difference in the hydrodynamic diameter versus the initial AuNPs-citrate (48 ± 32 nm vs 40 ± 25 nm). Remarkably, the LSPR band of the dissociated AuNPs was identical to the AuNPs-citrate: The color of the sample returned to its initial bright red color. This suggests that 100% of the aggregates were dissociated and that all AuNPs were detached and dispersed. For the remainder of this study, the efficiency of the dissociation will be characterized by the percentage of dissociation (%) as measured by UV-Vis spectroscopy (comparison between the LSPR band after dissociation versus that from AuNPs-citrate). See the experimental section in the supporting information for more details. Importantly, a concentration of 12 μM corresponds to a density of ∼4 HS-PEG_6_-OCH_3_/nm^2^. This finding is particularly interesting because the typical grafting density of HS-PEG_6_-OCH_3_ on dispersed AuNPs is reported to be between 3.5 and 4 HS-PEG_6_-OCH_3_ /nm^2^.^33^ This implies that no excess of HS-PEGs was needed to dissociate the entire assembly—only enough to cover all the gold surface. It is worth noting that the kinetics of the dissociation was almost instantaneous (equilibrium reached within 10 minutes) (Figure S6). A video of the dissociation of the AuNPs assemblies is available online.

These findings suggest that HS-PEG_6_-OCH_3_ can penetrate the assembly and graft onto the AuNPs surface, thus displacing the citrate and arginine layer. The particle becomes sterically stabilized and water-soluble when a sufficient amount of HS-PEG_6_-OCH_3_ is grafted on the particle surface; thus, the particle detaches from the assembly. When the concentration of HS-PEG_6_-OCH_3_ is sufficiently high to cover all of the particle surface, all assemblies dissociate, and the optical properties of the AuNPs are restored to those of the initial dispersed AuNPs-citrate. This phenomenon is possible only because (i) the AuNPs are not trapped in permanent aggregated state and (ii) HS-PEG_6_-OCH_3_ can replace the citrate/arginine layers due to the covalent grafting onto the gold surface.

To demonstrate the necessity of these two features, multiple control experiments were conducted. First, peptide-free conditions were used to promote the assembly of AuNPs-citrate and its dissociation with HS-PEG_6_-OCH_3_ was evaluated (**Figure 3A**). Here, salts were added to AuNPs-citrate to induce aggregation via disruption of the electrostatic repulsion forces between the particles. Even the addition of a high concentration of HS-PEG_6_-OCH_3_ (>100 μM) could not dissociate the assemblies and the color of the samples remained dull blue/grey. Unlike the peptide Arg-Arg, these ions cannot sterically prevent the AuNPs from falling in an irreversible aggregated state.

**Figure 3.**
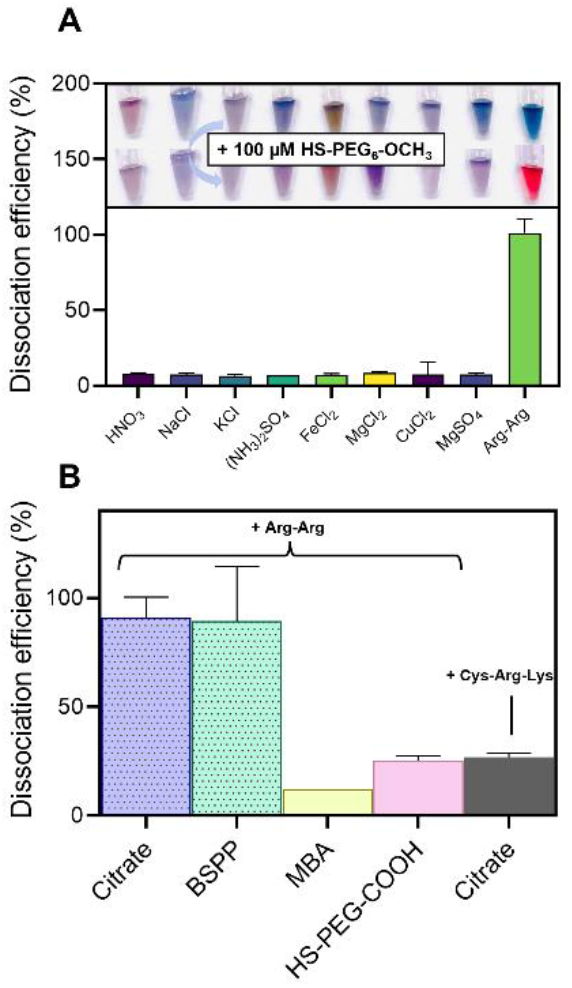
Dissociation efficiency of HS-PEG_6_-OCH_3_ for alternative assembly conditions or coating ligands. **(A)** Dissociation efficiency of 100 μM of HS-PEG_6_-OCH_3_ added to AuNPs-citrate assembled in peptide-free conditions. **(B)** Dissociation efficiency of 100 μM of HS-PEG_6_-OCH_3_ added to either AuNPs-citrate, AuNPs-BSPP, AuNPs-MBA, or AuNPs-S-PEG-COOH assembled with 10 μM of Arg-Arg or AuNPs-citrate assembled with 10 μM of Cys-Arg-Lys peptide.

AuNPs with different coating ligands were also evaluated and compared to citrate: bis(p-sulfonatophenyl)phenylphosphine (BSPP), mercaptobenzoic acid (MBA), or thiolated PEG-COOH (HS-PEG-COOH; Mw = 634 g.mol^−1^). Similar to the AuNPs-citrate, the presence of Arg-Arg led to the assembly of AuNPs-BSPP, AuNPs-MBA, and AuNPs-S-PEG-COOH. However, only the AuNPs-BSPP could be dissociated; assemblies of AuNPs-MBA or AuNPs-S-PEG-COOH were irreversible even in the presence of high concentrations (>100 μM) of HS-PEG_6_-OCH_3_ (**Figure 3B**). This is because MBA and HS-PEG-COOH make covalent Au-S bonds with the gold surface that prevent subsequent grafting of HS-PEG_6_-OCH_3_. In contrast, BSPP, like citrate, is only physiosorbed onto the gold surface and can be easily displaced by HS-PEG_6_-OCH_3_, thus making the assembly reversible.

Finally, an alternative control peptide was used for assembly (Cys-Arg-Lys). This peptide could aggregate the AuNPs-citrate but the assembly was irreversible (**Figure 3B**). This is because this peptide can make a covalent Au-S bond with the gold surface via the cysteine, thus hindering the grafting of HS-PEG_6_-OCH_3_ on the AuNPs surface and preventing dissociation. These control experiments show that the coating ligands as well as the aggregation peptide need to be weakly adsorbed onto the gold surface to facilitate aggregation and dissociation. Importantly, as the aggregates do not degrade over time: their dissociation was still possible even 120 days after their formation without a significant difference in the percent dissociation (Figure S7).

### Effect of the PEGs structure on the dissociation capacity

The impact of PEGs structure on dissociation capacity was studied next, i.e., ligands differing either by their size, anchoring group, core, or charge (**Table 1)**.

**Table 1.** Summary the ligands used to dissociate the AuNPs assemblies.

| Description | Name | Mw<br>(g.mol<sup>-1</sup>) | Anchor | Core | End group | Dissociation |
| --- | --- | --- | --- | --- | --- | --- |
| Effect of the<br>anchoring group | OH-PEG<sub>6</sub>-OCH<sub>3</sub> | 296 | OH | PEG<sub>6</sub> | Methoxy | 0 % |
|  | Alk-PEG<sub>4</sub>-OCH<sub>3</sub> | 232 | Alkyne | PEG<sub>4</sub> | Methoxy | 87 % |
|  | LA-PEG<sub>22</sub>-OCH<sub>3</sub> | 1000 | Lipoic acid | PEG<sub>22</sub> | Methoxy | 92 % |
| Effect of the<br>PEG core | MBA | 154 | Thiol | Benzene | Carboxyl | 0 % |
|  | MPA | 106 | Thiol | Propane | Carboxyl | 0 % |
| Effect of the<br>charge | HS-PEG<sub>226</sub>-NH<sub>2</sub> | 10,000 | Thiol | PEG<sub>226</sub> | Amine | 30 % |
| Effect of the<br>PEG size | HS-PEG<sub>450</sub>-OCH<sub>3</sub> | 20,000 | Thiol | PEG<sub>450</sub> | Methoxy | 20 % |
|  | HS-PEG<sub>226</sub>-OCH<sub>3</sub> | 10,000 | Thiol | PEG<sub>226</sub> | Methoxy | 35 % |
|  | HS-PEG<sub>110</sub>-OCH<sub>3</sub> | 5,000 | Thiol | PEG<sub>110</sub> | Methoxy | 60 % |
|  | HS-PEG<sub>44</sub>-OCH<sub>3</sub> | 2,000 | Thiol | PEG<sub>44</sub> | Methoxy | 75 % |
|  | HS-PEG<sub>22</sub>-OCH<sub>3</sub> | 1,000 | Thiol | PEG<sub>22</sub> | Methoxy | 91 % |
|  | HS-PEG<sub>6</sub>-OCH<sub>3</sub> | 356 | Thiol | PEG<sub>6</sub> | Methoxy | 100 % |
|  | HS-PEG<sub>4</sub>-OCH<sub>3</sub> | 224 | Thiol | PEG<sub>4</sub> | Methoxy | 100 % |

First, to confirm that the grafting of the PEG on the AuNPs is critical to dissociation, methoxy PEG molecules carrying different anchoring groups (x-PEG-OCH_3_) were studied. **Figure 4A** shows that the dissociation of Arg-Arg-AuNPs assemblies is no longer possible when the thiol group (HS-PEG_4_-OCH_3_) is replaced by a hydroxy group (OH-PEG_4_-OCH_3_) even at high concentrations (>100 μM): This is because OH-PEG_4_-OCH_3_ cannot bind covalently to the gold surface. Positive controls involving alkyne (Alk-PEG_4_-OCH_3_) and lipoic acid (LA-PEG_22_-OCH_3_) as anchoring groups were investigated because these two chemical groups, like thiol groups, can form a covalent bond with the gold atoms at the AuNPs surface.^34,35^ Thus, as expected, the addition of Alk-PEG_4_-OCH_3_ or LA-PEG_22_-OCH_3_ led to the dissociation of the AuNPs assemblies (**Figure 4B**). Interestingly, while LA-PEG_22_-OCH_3_ had a similar dissociation efficiency than its thiolated counterpart (HS-PEG_22_-OCH_3_), the one of Alk-PEG_4_-OCH_3_ was lower compared to HS-PEG_4_-OCH_3_. It can be explained by the fact that alkynes form a less labile strong bond with the surface compared to thiols. Overall, the results confirm the necessity for the dissociating ligands to be capable of grafting onto the AuNPs surface.

**Figure 4.**
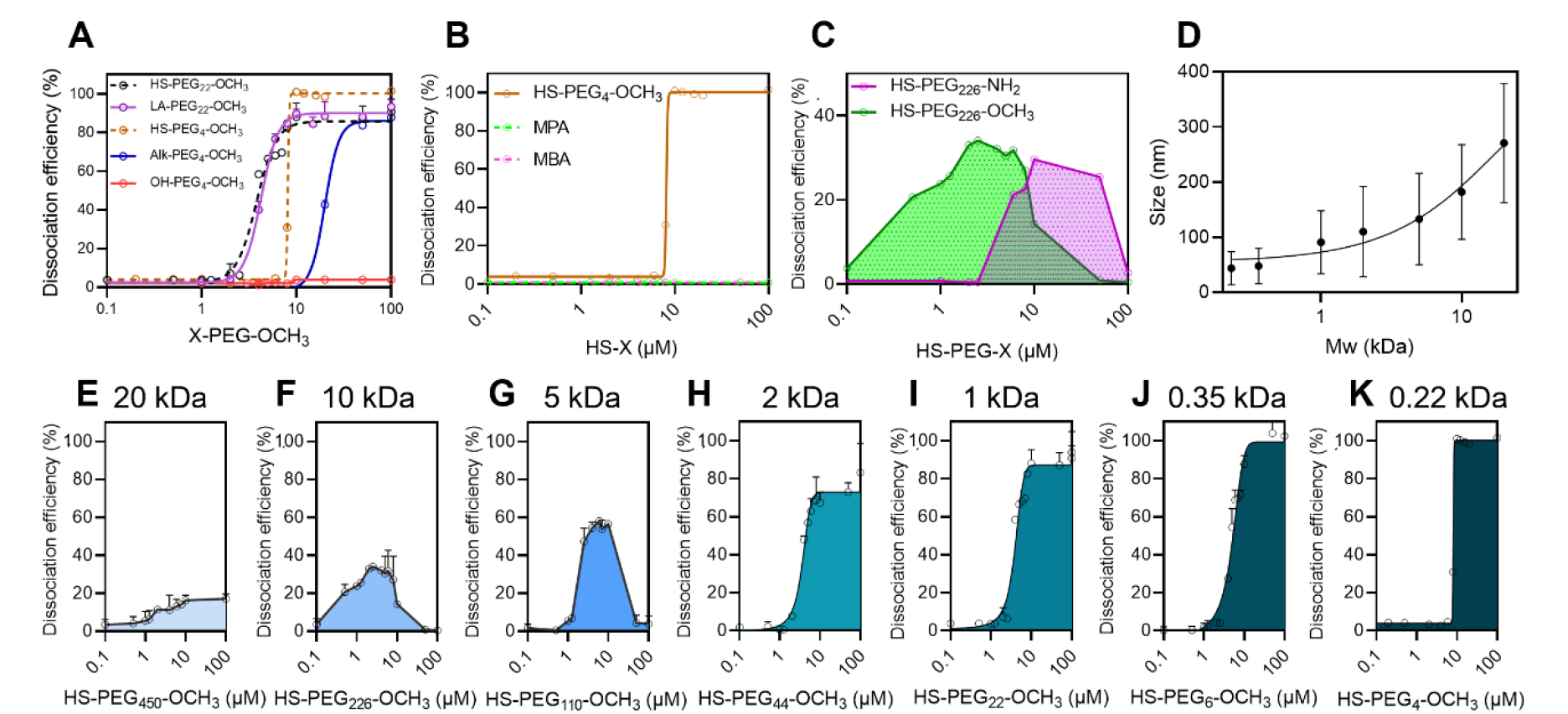
Effect of the PEG properties on the dissociation efficiency. **(A)** Dissociation efficiency of X-PEG-OCH_3_ molecules carrying various anchoring groups added to Arg-Arg-AuNPs. X= thiol (HS), hydroxy (OH), lipoic acid (LA) or Alkyne (Alk). **(B)** Dissociation efficiency of HS-PEG_4_-OCH_3_, MBA, or MPA added to Arg-Arg-AuNPs. **(C)** Dissociation efficiency of HS-PEG_226_-OCH_3_ compared to HS-PEG_226_-NH_2_ added to Arg-Arg-AuNPs. **(D)** Size of the assembly 20 minutes after the addition of HS-PEG_x_-OCH_3_ to Arg-Arg-AuNPs of different molecular weight (20, 10, 5, 2, 1, 0.35 and 0.22 kDa). **(E), (F), (G), (H), (I), (J)** and **(K)**. Dissociation efficiency of HS-PEG_x_-OCH_3_ concentration with different molecular weight added to Arg-Arg-AuNPs.

Non-PEG ligands such as mercaptobenzoic acid (MBA) and mercaptopropionic acid (MPA) were investigated next: Interestingly, none could dissociate the aggregates despite the presence of a thiol group in their structures (**Figure 4B**). The flexibility and hydrophilicity of the PEG chain is likely crucial to make the AuNPs water-soluble and detach them from the bulky and hydrophobic aggregate.

Finally, we demonstrated that the charge and the size of the PEG molecules have a strong impact on their capacity to dissociate AuNPs aggregates. **Figure 4C** shows that the dissociation of HS-PEG_226_-OCH_3_ is strongly affected when the methoxy group is replaced by an amine group (net charge = +1). The effect of the size of the PEG molecule was then studied by using various HS-PEGx-OCH_3_ of different molecular weights ranging from 20 kDa to 0.22 kDa. **Figure 4D** shows that the size of the assemblies (measured by MANTA) is inversely proportional to the size of the HS-PEGx-OCH_3_. This result was confirmed by UV-Vis spectroscopy: HS-PEG_x_-OCH_3_ with a molecular weight of 20, 10, 5 and 2 kDa, could only dissociate a maximum of 20%, 35%, 60%, and 70% of the AuNPs assemblies, respectively (**Figures 4E, 4F, 4G** and **4H**). However, the 1 kDa HS-PEG_x_-OCH_3_ offered 80% dissociation; this value reached 100% for HS-PEG_x_-OCH_3_ with a molecular weight of 0.35 kDa or smaller (**Figures 4I, 4J** and **4K**). These data illustrate that the dissociation efficiency of PEG is directly proportional to PEG size. Large HS-PEGs likely cannot reach the interfaces of the aggregated particles while smaller HS-PEGs can. However, the concentration of HS-PEG_x_-OCH_3_ needed to initiate the dissociation was proportional to their size because the footprint of HS-PEGs is directly dependent on their size. Thus, larger HS-PEGs cover a larger surface on the AuNPs and less PEGs molecules are needed to complete the coating. This non-exhaustive study reveals that the dissociation of AuNPs assemblies can be directly modulated by tuning the PEGs properties, and it is particularly interesting for the development of sensing strategies.

### Advantages of the dissociation approach

Our dissociation approach possesses two major advantages compared to the conventional aggregation-based assay. First, it can operate across various complex samples that is crucial for assay generalizability. Typically, background interferents differ across different sample matrices; thus, it is challenging to obtain an unambiguous colorimetric signal that is insensitive to the matrix composition—particularly for AuNPs aggregation-based assays because the colloids can be unstable in these conditions or because endogenous molecules can prevent the aggregation. The dissociation of the AuNPs assembly is very robust: It is unaffected by high ionic strength (>1 M NaCl) (Figure S8) or pH extremes (3 or 13) (Figure S9). Only compounds that can interact with the thiol group can interfere with the dissociation. As an example, dithiothreitol (DTT) quenching is presented in Figure S10. Thus, we next evaluated dissociation as a function of sample type.

The dissociation of Arg-Arg-AuNPs assemblies was investigated in pooled plasma, pooled urine, pooled saliva, pooled bile, human embryonic kidney (HEK) 293 cell lysates in Dulbecco’s modified Eagle’s medium (DMEM), and even sea water. Importantly, the matrices by themselves could not dissociate the assemblies even after three hours of incubation (Figure S11). This is particularly interesting because these complex environments usually produce dramatic background signals. For example, the dispersibility of AuNPs-citrate is strongly impacted when suspended in urine or sea water due to the high ionic strength (Figure S12). The dissociation of Arg-Arg-AuNPs in complex matrices was investigated with 10 μM HS-PEG_6_-OCH_3_. Typically, concentrated Arg-Arg-AuNPs were suspended in the complex matrix and then HS-PEG_6_-OCH_3_ was added. The complex matrix represented at least 90% of the total volume. At least 80% of dissociation was obtained in all the matrices including an unambiguous color change from blue to red (**Figure 5A** and **5B**).

**Figure 5.**
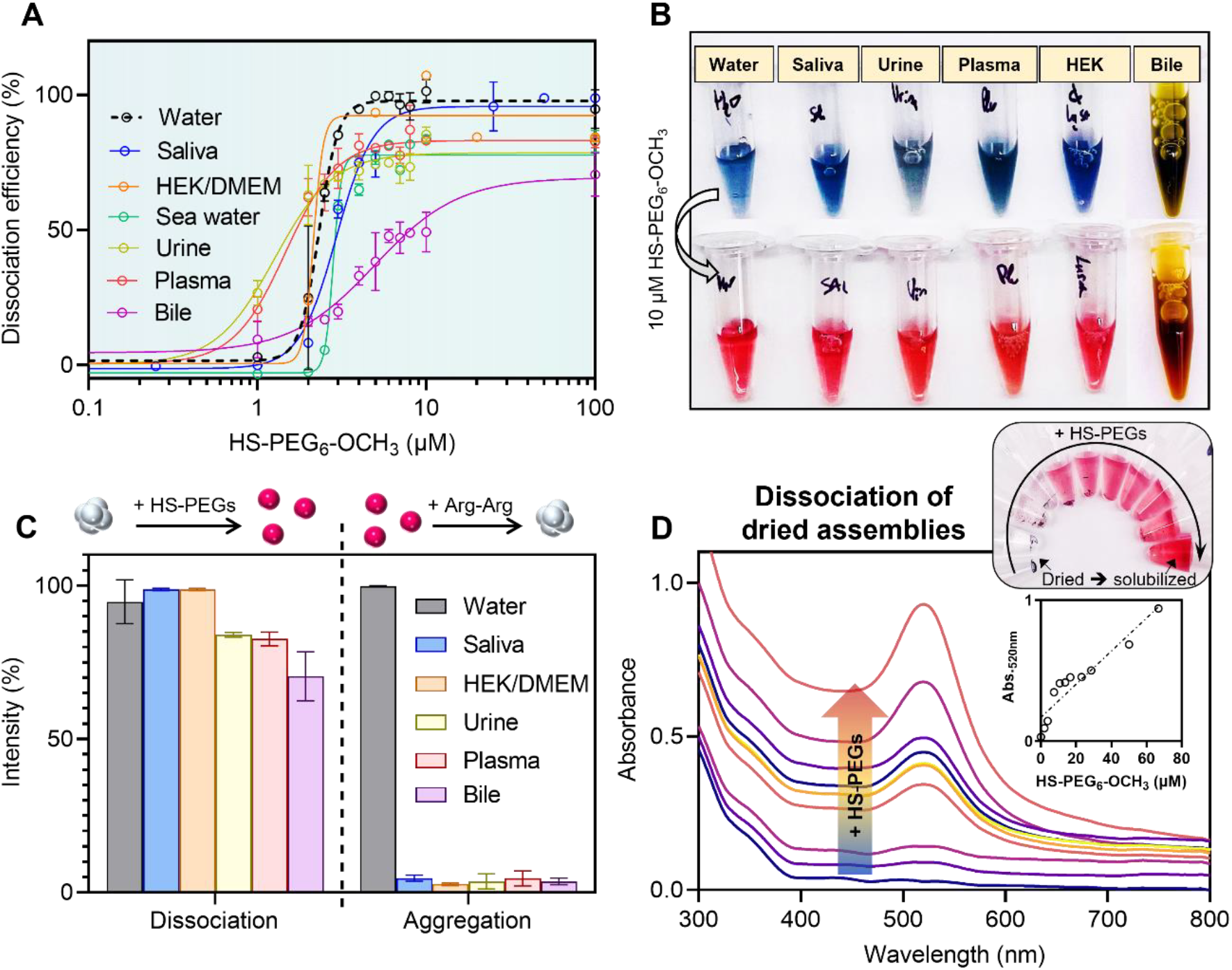
Dissociation of AuNPs assemblies in complex matrices. **(A)** Dissociation efficiency of HS-PEG_6_-OCH_3_ added to Arg-Arg-AuNPs suspended in various media and **(B)** the corresponding pictures before and 20 minutes after the addition of 10 μM of HS-PEG_6_-OCH_3_. The complex medium was approximately 90% of the total volume except for the bile that was 20%. **(C)** Dissociation of Arg-Arg-AuNPs with HS-PEG_6_-OCH_3_ compared to the Arg-Arg-induced aggregation of AuNPs-citrate in various complex matrices. Note that HEK/DMEM corresponds to HEK 293 cell lysates coming from suspension of 10^6^ cells in Dulbecco’s modified Eagle’s media with 10% fetal bovine serum and 1% penicillin-streptomycin. **(D)** UV-Vis spectra of dried Arg-Arg-AuNPs film after dissociation with aqueous solution of HS-PEG_6_-OCH_3_ of various of concentrations. Insets show the absorbance at 520 nm as a function of the concentration of HS-PEG_6_-OCH_3_ and the corresponding pictures.

The re-dispersion of AuNPs assemblies was then compared to the aggregation of dispersed AuNPs-citrate with the Arg-Arg peptide in complex media (**Figure 5C**). Even high concentrations of Arg-Arg (>200 μM) could not induce the AuNPs-citrate aggregation in biological fluids because of the numerous endogenous molecules/proteins that shield the electrostatic interactions between the peptide and the particles as well as the formation of a protein corona around the particles.^17^ The matrix-insensitive feature of our dissociation design is explained by the fact that most of the biomolecules found in the complex samples (proteins, phospholipids, nucleotides, etc.) can only stick onto the gold surface via van der Waals forces, electrostatic or hydrophobic interactions, or hydrogen bonds.^17,36^ Thus, the grafting of HS-PEGs is favored because the Au-S bond energy is approximately 10-fold higher than the average hydrogen bond.^37^ Also, PEGs are commonly used to prevent non-specific adsorption of proteins on AuNPs, and thus the HS-PEGs do not interact much with the interferents; they remain free to bind to the particles.

The second advantage of our dissociation approach is that the assemblies did not just remain aggregated—they also precipitated, thus increasing the color change between the dissociated and aggregated systems. As an example, Figure S13 shows that Arg-Arg-AuNPs suspended in saliva after 2 hour tend to precipitate, thus leading to a loss of blue color in the suspension while the dissociated particles remain bright red. Interestingly, even if precipitated, the assemblies can still be dissociated with 10 μM of HS-PEG_6_-OCH_3_.

Remarkably, the HS-PEGs could even solubilize dried films of Arg-Arg-AuNPs (**Figure 5D**). Briefly, 50 μL of aqueous solution containing different concentrations of HS-PEGs, ranging from 0 to 70 μM, were added to the dried particles and 20 minutes later, the UV-Vis spectrum of the solution was recorded. In the absence of HS-PEGs, no LSPR band of the particles was observed as no particles detached spontaneously and the solution remained clear. However, in the presence of HS-PEGs, the plasmonic band of the particles was observed proportionally to the HS-PEGs concentration without the need of sonication. The HS-PEGs led to the spontaneous and progressive detachment of the AuNPs from the surface and thus a color change (**Figure 5D, inset**). The solubilization of the AuNPs could be thus quantified using only the absorbance at 520nm in the solution as the LSPR band intensity was linearly proportional to the HS-PEGs concentration. This approach did not work on AuNPs-citrate as they degraded during the drying process—the di-peptide is a crucial additive. Finally, the solubilization of dried Arg-Arg-AuNPs was possible even in biofluids such as saliva, plasma, urine or cell lysates (see supporting information Figure S14).

Thus, our dissociation strategy affords two advantages versus traditional aggregation-based assays: It is insensitive to the composition of the operating medium and the gap of colorimetric signals between dissociated and non-dissociated samples is unambiguous and increases over time, thus enhancing the naked eyes identification.

### Protease detection

The charge and size of the HS-PEGs modulate their capacity to dissociate the AuNPs assemblies, and thus we designed a strategy to detect proteases in biological fluids. Trypsin was chosen as a model protease because it possesses a high catalytic efficiency and easily cleaves peptide sequences after arginine or lysine.^38^ Trypsin is a biomarker of pancreatic cancer and can be found in micromolar concentrations in blood or urine.^39^

Peptides containing the motif Arg-Arg-Lys (RRK) were thus conjugated to a thiolated PEG capped with a carboxyl group (HS-PEG_12_-COOH, Mw = 634 Da) via EDC/NHS cross-linking reaction. Three peptides based on the motif RRK were investigated and the number of RRK repetitions varied from 1 to 3 to increase the mass and charge of the resulting HS-PEG-peptide conjugate (**Figure 6A**).

**Figure 6.**
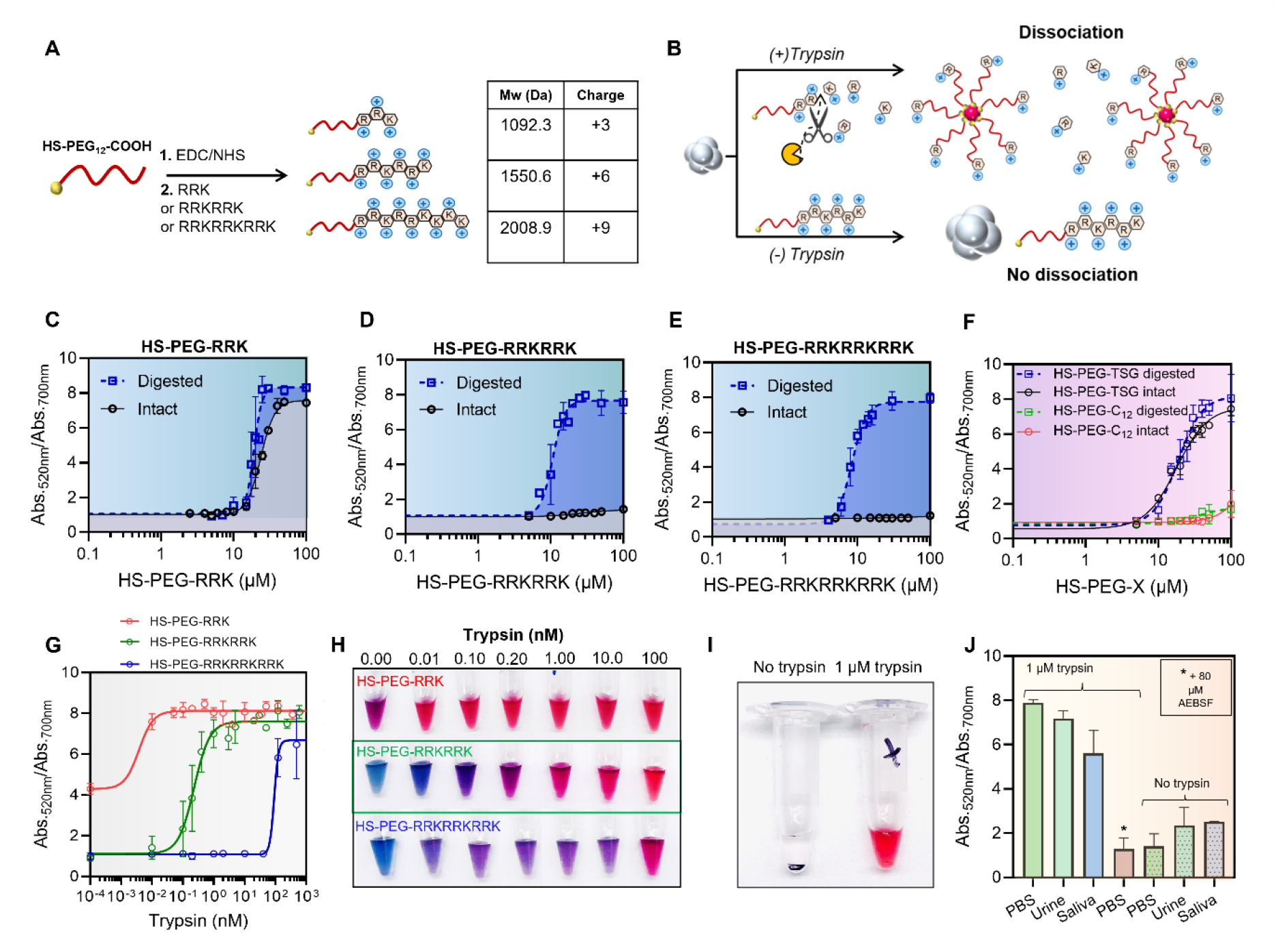
Trypsin sensing with HS-PEG-peptide conjugates. **(A)** Scheme of the synthesis of HS-PEG-peptide conjugates via EDC/NHS chemistry including the three main sequences investigated as well as the size and charge of the compounds. **(B)** Illustration of the sensing mechanism: The trypsin cleavage reduces the size and charge of the HS-PEG-peptide conjugate, which increases their capacity to dissociate the AuNPs assemblies. Dissociation of Arg-Arg-AuNPs assemblies with either **(C)** HS-PEG-RRK, **(D)** HS-PEG-RRKRRK, or **(E)** HS-PEG-RRKRRKRRK intact (black line) or digested by 1 μM of trypsin for 24 h at 37 °C (dashed blue line). **(F)** Dissociation of Arg-Arg-AuNPs assemblies with HS-PEG-TSG and HS-PEG-C_12_ either intact or digested by trypsin. **(G)** Trypsin detection using the three different HS-PEG-peptide conjugates and **(H)** the corresponding pictures. **(I)** Picture of dried Arg-Arg-AuNPs assemblies after the addition of 50 μL of HS-PEG-RRKRRK incubated or not with 1 μM of trypsin. **(J)** Comparison between the detection of 1μM of trypsin spiked either in PBS or in urine as well as in PBS with 80 μM of AEBSF inhibitor.

From concentrations of 10 μM, HS-PEG_12_-COOH is capable to dissociate 100% of the assemblies (Figure S15). However, the conjugation of the peptides to the HS-PEG_12_-COOH was expected to decrease its capacity to dissociate the assemblies as it increases its size and charge. In this context, the addition of the HS-PEG-peptide conjugates to AuNPs assemblies could not generate a colorimetric signal. However, in the presence of trypsin, the proteolytic cleavage could progressively remove the amino acids residues from the conjugate, thus reducing its size and charge that would restore the dissociation capacity. A colorimetric signal, proportional to the trypsin activity, could thus be observed (**Figure 6B**).

The HS-PEG-peptide conjugates were synthesized and titrated to Arg-Arg-AuNPs assemblies in order to evaluate the quenching of their dissociation capacity. EDC/NHS cross-linking between HS-PEG-COOH and RRKRRK or RRKRRKRRK led to a total quenching of the dissociation capacity of HS-PEG_12_-COOH even at high concentrations (>100 μM) (**Figures 6D** and **6F**) while the conjugation of RRK only reduced it slightly (**Figure 6E**). In the absence of EDC/NHS, no cross-linking could occur and the dissociation capacity of HS-PEG_12_-COOH was not affected (Figure S16).

Subsequently, the three conjugates were digested with trypsin (10 μM, 37°C, 24h) and titrated into the AuNPs assemblies again. Interestingly, the dissociation capacity of HS-PEG-RRKRRK and HS-PEG-RRKRRKRRK was restored; HS-PEG-RRK was also enhanced because cleavage by trypsin reduced the size and the positive charge of the conjugate (**Figure 6C, 6D** and **6E**). A minimum of two repetitions of the motif RRK is thus necessary to ensure a total quenching of the dissociation capacity of the HS-PEG_12_-COOH. This can later be completely restored with the proteolytic cleavage. Only one repetition of RRK leads to a very narrow window of detection between the intact and digested HS-PEG-peptide. In addition to conferring a higher positive charge, the RRKRRK and RRKRRKRRK peptides could conjugate multiple HS-PEGs per peptide due to the presence of lysine residues that in turn lead to an even bigger construct. Matrix-assisted laser desorption/ionization (MALDI) was used to characterize the HS-PEG-RRKRRK conjugate (Figure S17) and a mixture of single and multiple conjugations of HS-PEG per peptide were observed.

Control experiments were then conducted with alternative non-trypsin cleavable molecules: the peptide TSG and the bis-amine C_12_. The resulting conjugates (HS-PEG-TSG and HS-PEG-C_12_) were then titrated to Arg-Arg-AuNPs assemblies before and after digestion with trypsin, similarly to what was described previously. The dissociation capacity of HS-PEG-TSG was similar to the one of HS-PEG_12_-COOH because TSG is a short peptide and lacks positive charge. It is not impacted by the digestion by trypsin. On the other hand, the conjugation of C_12_ led to total quenching of the dissociation capacity that could not be restored with trypsin digestion. This is because C_12_, like RRKRRK and RRKRRKRRK, contains more than one free amine and can thus be conjugated to multiple HS-PEG molecules. This makes the conjugate too bulky to dissociate the AuNPs. However, unlike RRKRRK and RRKRRKRRK, C_12_ does not have any cleavable site for trypsin and thus the trypsin digestion has no effect on the dissociation capacity (**Figure 6F**).

## CONCLUSION

In summary, the dissociation of AuNPs assemblies with HS-PEGs molecules was studied and exploited to build matrix-insensitive sensors. Robust assemblies of AuNPs-citrate were formed using a di-arginine additive (Arg-Arg). The efficient electrostatic interactions between the citrate and the arginine led to compact assemblies of the particles, thus provoking a strong modification of their optical properties. However, the presence of peptides protected the AuNPs from degradation. Surprisingly, the addition of HS-PEGs could dissociate the assemblies with a total recovery of the initial optical properties. The mechanism was fully characterized by TEM, MANTA, UV-Vis, and FTIR spectroscopies. The HS-PEGs can progressively graft onto the AuNPs surface and remove the citrate/arginine layers. As the hydrophilic PEG layer surrounds the AuNPs, the particles progressively detach from the bulky assemblies and become water-dispersible. Importantly, only a minimum amount of HS-PEGs is needed to cover all the gold surfaces (∼ 4 HS-PEG_6_-OCH_3_/nm^2^). We have thus shown that the dissociation capacity of HS-PEGs is modulated by their size and charge. HS-PEG-OCH_3_ with a molecular weight of 1000 Da or less could dissociate 80% or more. Remarkably, the dissociation of the assemblies was matrix-insensitive and produced an unambiguous color change in plasma, saliva, urine, bile, cell lysates, or even sea water. Moreover, we found that the generation of the colorimetric signal could be improved by using dried film of AuNPs assemblies. The presence of HS-PEGs leads to the detachment of AuNPs from the surface as it solubilizes them. The color of the suspension becomes then red and its intensity is proportional to the amount of HS-PEGs. In the absence of this later, the color of the solution is clear. This strategy allows a unambiguous distinction with the naked eyes between samples that have or not HS-PEGs. We thus designed a sensing strategy based on HS-PEG-peptide probes and AuNPs assemblies as signal read out for protease sensing in complex media. Trypsin was chosen as model protease and peptide containing repetition of the motif RRK were conjugated to HS-PEGs. The optimized conjugate, HS-PEG-RRKRRK, allowed the visual detection of trypsin with a picomolar limit of detection. Detection could be performed simply in pooled urine or saliva spiked with trypsin. To the best of our knowledge, this is the first time that HS-PEGs molecules have been used to dissociate AuNPs assemblies or to solubilize dried AuNPs and combined with peptides for protease detection. This innovative approach could benefit protease detection across various complex environments. The approach could be adapted to any protease as long as a peptide substrate can be conjugated to HS-PEGs and the dissociation capacity of the resulting conjugate can only be restored by the proteolytic activity.

## ASSOCIATED CONTENT

### Supporting Information

Materials, instrumentations, methods, video of AuNPs dissociation, TEM images, stability over time, dissociation kinetics, dissociation in NaCl, dissociation at different pH, and dissociation with DTT, 40 nm-NPs study.

### Notes

The authors declare no competing financial interest.

## ACKNOWLEDGMENT

The authors thank internal funding from the UC Office of the President (R00RG2515) and the National Institutes of Health (R01 DE031114; R21 AG065776; R21 AI157957; 1 S10 OD023555-01A1) for financial support. This work was supported in part by the National Science Foundation Graduate Research Fellowship Program under Grant No. DGE-1650112. The electron microscopy work was performed in part at the San Diego Nanotechnology Infrastructure (SDNI) of University of California San Diego, a member of the National Nanotechnology Coordinated Infrastructure (NNCI), which is supported by the National Science Foundation (Grant ECCS-1542148). The MANTA analysis work was supported by the National Institutes of Health (S10 OD023555). MR thanks the Wallonie-Bruxelles International Foundation. This work used equipment purchased supported by the UC San Diego Materials Research Science and Engineering Center (UCSD MRSEC), supported by the National Science Foundation (Grant DMR-2011924).

