## SUpporting in for "A di-arginine additive for dissociation of gold nanoparticle aggregates: A matrix-insensitive approach with applications in protease detection"

### 1. Materials

Bis(*p*-sulfonatophenyl)phenylphosphine dihydrate dipotassium salt (BSPP, 97%), gold(III) chloride trihydrate ($HAuCl_4{\cdot}3H_2O$, >99.9%), sodium citrate tribasic dihydrate (>99%), trypsin, 4-mercaptobenzoic acid (MBA, 99%), 3-mercaptopropionic acid (MPA, >99%), N-hydroxysuccinimide (NHS, 98%), 1-12-diaminododecane (C12, 98%), O-(2-mercaptoethyl)-O′-methyl-hexa(ethylene glycol) (HS-$PEG_6$-$OCH_3$, 98%), DL-dithiothreitol (DTT, >99%) and pooled human plasma were purchased from Sigma Aldrich (St. Louis, MO). HS-PEG-$OCH_3$ Mw = 20,000, 10,000, 5,000 and 2,000 g.mol$^{-1}$ and HS-PEG-$NH_2$ Mw = 10,000 g.mol$^{-1}$ were purchased from Laysan Bio, Inc. (Arab, AL). HS-PEG-$OCH_3$ Mw = 1,000 g.mol$^{-1}$ and LA-PEG-$OCH^3$ Mw = 1000 g.mol$^{-1}$ was purchased from Biopharma PEG (Watertown, MA). HS-PEG-$OCH_3$ Mw = 224 g.mol$^{-1}$, HS-PEG-COOH Mw = 600 g.mol$^{-1}$, Alk-PEG-OH Mw = 232.27 g.mol$^{-1}$, and OH-PEG-$OCH_3$ Mw = 296.36 g.mol$^{-1}$ were purchased from PurePEG (San Diego, CA). AuNPs-citrate (40 nm) were purchased from Nanocomposix (San Diego, CA). The (1-ethyl-3-(3-dimethylaminopropyl)carbodiimide hydrochloride) (EDC) was purchased from Thermofisher (Waltham, MA). Gibco Phosphate Buffer Saline (PBS) pH 7.4 was purchased from Fisher Scientific (Pittsburgh, PA). The 4-(2-Aminoethyl)-benzenesulfonyl fluoride, HCl (AEBSF) was purchased from Research Product International (Mount Prospect, IL). Peptides Arg-Arg-Lys (RRK), Arg-Arg-Lys-Arg-Arg-Lys (RRKRRK) and Arg-Arg-Lys-Arg-Arg-Lys-Arg-Arg-Lys (RRKRRKRRK) were purchased from Genscript Probio (Piscataway, NJ). Sodium chloride (NaCl), potassium chloride (KCl), ferrous chloride ($FeCl_2$), nitric acid ($HNO_3$), copper (II) chloride ($CuCl_2$), and magnesium chloride ($MgCl_2$) were purchased from Fisher Scientific International, Inc. (Hampton, NH). Pooled human saliva was purchased from Lee Biosolutions, Inc. (Maryland Heights, MO). Pooled human urine filtered was purchased from Innovative Research (Novi, MI). Human bile was purchased from Zen Bio (Durham, NC). Sea water was collected in Pacific Beach (San Diego, CA). Fmoc-protected L/D-amino acids, hexafluorophosphate benzotriazole tetramethyl uronium (HBTU), and Fmoc-Rink amide MBHA resin (0.67 mmol/g, 100-150 mesh) were purchased from AappTec, LLC (Louisville, KY). Organic solvents including N,N-dimethylformamide (DMF, sequencing grade), acetonitrile (ACN, HPLC grade), ethyl ether (certified ACS), methylene chloride (DCM, certified ACS), and dimethyl sulfoxide (DMSO, certified ACS) were from Fisher Scientific International, Inc. (Hampton, NH). Ultrapure water (18 MΩ·cm) was obtained from a Milli-Q Academic water purification system (Millipore Corp., Billerica, MA). Amicon® ultra-15 centrifugal filter units (M.W. cutoff =100 kDa) and automation compatible syringe filters (PTFE, 0.45 mm) were from MilliporeSigma (St. Louis, MO). Glassware and stir bars were cleaned with aqua regia ($HCl$:$HNO_3$=3:1 by volume) and boiling water before use. Beside aqua regia that has to be handled carefully, no unexpected or unusually high safety hazards were encountered.

### 2. Instrumentation and methods

#### 2.1 AuNPs synthesis

Citrate-stabilized AuNPs (~20 nm) were prepared using the Turkevich method by rapidly injecting an aqueous solution of sodium citrate tribasic dihydrate (150 mg, 5 mL) into an aqueous solution of $HAuCl_4.3H_2O$ (45 mg, 300 mL) under boiling conditions and vigorous stirring. The reaction mixture was left boiling while stirring for another 15 min and then cooled down to room temperature. The deep red dispersion was then purified by applying one round of centrifugation at 18,000 g for 30 min, and the pink supernatant was discarded. The resulting pellet of AuNPs-citrate was redispersed in deionized water by sonication and stored at ambient conditions.

#### 2.2 AuNPs assembly and dissociation

**Assembly.** Typically, 1 mL of AuNPs-citrate at OD = 1.5 was mixed with 50 µL of Arg-Arg (100 µM) to provoke the AuNPs assembly. The color of the suspensions rapidly changed from red to blue. The assembled AuNPs (Arg-Arg-AuNPs) were stable over time when stored at 4°C and could be used for dissociation even month after their assembly. **Dissociation.** Stock solutions of the different HS-PEGs with concentrations ranging from 10 µM to 1 mM were prepared. Specific volumes of each solution were added to a 96-well plate in order to reach the desired final HS-PEGs concentration and then 100 µL of Arg-Arg-AuNPs were added. For complex matrices experiments, the Arg-Arg-AuNPs were first concentrated 10 times by centrifugation and then dispersed in complex media (sea water, pooled human saliva, plasma, urine, bile or HEK cell lysates) that represented thus 90% of the total volume except for bile, which was only 20%. Quickly after the addition of AuNPs, the dissociation of the assembly was monitored and the ratio of the absorbances at 520 nm and 700nm was recorded over time. The percentage of dissociation is described as follows:

$$\% \text{ of dissociation} = \frac{\left(\frac{Abs520nm}{Abs700nm}\right)_{AuNPs-S-PEGs} - \left(\frac{Abs520nm}{Abs700nm}\right)_{Arg-Arg-AuNPs}}{\left(\frac{Abs520nm}{Abs700nm}\right)_{AuNPs-citrate} - \left(\frac{Abs520nm}{Abs700nm}\right)_{Arg-Arg-AuNPs}}$$

Where $\left(\frac{Abs520nm}{Abs700nm}\right)_{AuNPs-S-PEGs}$ is the ratio of the absorbance after dissociation with HS-PEGs, $\left(\frac{Abs520nm}{Abs700nm}\right)_{AuNPs-citrate}$ is the ratio of the absorbance of the initial citrate-capped AuNPs, and $\left(\frac{Abs520nm}{Abs700nm}\right)_{Arg-Arg-AuNPs}$ is the ratio of the arginine-induced assembly of AuNPs.

#### 2.3 Drying and solubilization of the AuNPs assemblies

First, 1 mL of AuNPs-citrate was concentrated 5 times by centrifugation (18,000g during 18 minutes). Then, the assembly was provoked by adding 25 µL of Arg-Arg (100 µM) to the resulting 200 µL of concentrated AuNPs-citrate. Finally, 10 µL of the concentrated Arg-Arg-AuNPs were added to an Eppendorf and dried at 40 °C overnight. The solubilization of the dried assemblies with HS-PEGs was performed similarly to the dissociation of the assemblies. It is worth noting that shaking vigorously the Eppendorf could be required.

#### 2.4 Peptide synthesis

Peptides Arg-Arg (RR), Arg-Arg-Arg-Arg-Arg (RRRRR), Arg-Gly-Gly-Gly-Arg (RGGGR) and Tyr-Ser-Gly (TSG) were synthesized using an automated Eclipse™ peptide synthesizer (AAPPTec, Louisville, KY) through standard solid phase Fmoc synthesis on Rink-amide resin. Peptides were lyophilized in a FreeZone Plus 2.5 freeze dry system (Labconco Corp., Kansas, MO).

Peptide purification used a Shimadzu LC-40 HPLC system equipped with a LC-40D solvent delivery module, photodiode array detector SPD-M40, and degassing unit DGU-403. The crude sample was dissolved in an acetonitrile/$H_2O$ mixture (1:1, v/v) with an injection volume of 2 mL. This was applied on a Zorbax 300 BS, C18 column (5 µm, 9.4 × 250 mm) from Agilent (Santa Clara, CA) and eluted at a flow rate of 1.5 mL/min over a 40 min linear gradient from 10% to 95% of acetonitrile in water (with 0.05% TFA, HPLC grade). Preparative injections were monitored at an absorbance of 190, 220, and 254 nm. Fractions containing the pure peptide as confirmed by electrospray ionization mass spectroscopy were lyophilized and aliquoted (see below). All peptides were purified by HPLC to reach a purity of at least 90%.

Peptide synthesis and cleavage was confirmed using Electrospray ionization mass spectrometry (ESI-MS, positive ion mode) via the Micromass Quattro Ultima mass spectrometer in the Molecular MS Facility (MMSF). ESI-MS samples were prepared in a MeOH/$H_2O$ mixture (1:1, v/v).

#### 2.5 HS-PEG-peptide conjugate synthesis

Typically, 1 mL of HS-$PEG_{12}$-COOH (HS-$(CH_2CH_2O)_{12}$-$CH_2CH_2$COOH, 634 Da, 1 mM) dissolved in MES buffer (10 mM, pH = 5.5) was activated via the addition of 100 µL of EDC (50 mM, MES) and 40 µL of NHS (500 mM, MES). The reaction was stirred at room temperature for one hour. After one hour, 1 mL of peptide (or $NH_2$-molecule) (1 mM) dissolved in PBS (100 mM, pH = 7.4) was added to the activated HS-$PEG_{12}$-COOH. The reaction was stirred for 4 hours at room temperature and then stored at 4 °C.

#### 2.6 Trypsin incubation

Typically, 5 µL of the PEG-peptide conjugate (0.44 mM) were added to 5 µL of PBS 1x. Subsequently, 2 µL of trypsin of various concentrations were added and the resulting solution was incubated at 37.5 °C for 2 hours. At the end of the incubation, 100 µL of aggregated AuNPs (Arg-Arg-AuNPs) were added, and a color change proportional to the trypsin concentration was observed. UV-Vis spectra were recorded 30 minutes later.

#### 2.7 Trypsin inhibition study

Aqueous AEBSF inhibitor solution (20 mM) was prepared, and a specific volume was mixed with 1 µM of trypsin in order to reach the desired final AEBSF concentration. The resulting mixture was stirred gently at room temperature for 30 minutes. Subsequently, the specific volume of HS-PEG-RRKRRK was added to reach a final concentration of 200 µM and the solution was incubated at 37.5 °C for 2 hours. Finally, 100 µL of Arg-Arg-AuNPs (20 nm) were added, and the UV-Vis spectrum was recorded 10 minutes later.

#### 2.8 UV-Vis spectroscopy

The optical absorption measurements were collected using a hybrid multi-mode microplate reader (Synergy™ H1 model, BioTek Instruments, Inc.) in a 96-well plate. The dissociation of assembly was characterized by measuring the ratio of the absorbance at 520 nm or 530 nm and 700 nm or 820 nm for 20 nm NPs or 40 nm NPs, respectively.

#### 2.9 ATR-FTIR spectroscopy

ATR-FTIR spectra were recorded with a Nicolet™ iS50 FTIR Spectrometer with a DLaTGS detector by natural drying of 1 µL of AuNPs suspensions. Typically, 1 mL of AuNPs (2 nM) was first cleaned from unbound molecules via four cycles of centrifugation (18,000g during 18 minutes) and resuspended in 40 µL of pure water. The final concentration was then approximately 50 nM.

#### 2.10 Transmission Electron Microscopy (TEM)

Transmission electron microscopy (TEM) images of the Au colloids were acquired using a JEOL 1200 EX II operating at 80 kV. The TEM grids were prepared by natural drying of 2 µL of 2 nM AuNPs.

#### 2.11 Multispectral Advanced Nanoparticles Tracking Analysis (MANTA)

The multispectral advanced nanoparticle tracking analysis (MANTA) is a technique that builds images from the particles' light scattering via three lasers of different wavelengths (e.g., blue, green, and red); the wavelength of scattering depends on nanoparticle size. Thus, small particles (<100 nm) appear blue while larger particles appear greener or redder. MANTA uses these images to count the nanoparticles and calculate their size based on Brownian motion. MANTA was performed with the ViewSizer 3000 (Horiba Scientific, USA). The temperature was set to 25 °C during the measurement. Automated noise analysis determines the optimal wavelength for representing each nanoparticle. Here, 8-bit composite videos were generated, and 10 videos were used per analysis (300 frames for seconds). A quartz cuvette with minimum volume of 1 mL was used for the measurement and the AuNPs concentrations was set up at 0.04 nM.

#### 2.12 Cell culture

A human embryonic kidney cell line (HEK 293T) was used for this work. The cells were cultured in Dulbecco's modified Eagle's medium (DMEM) with 10% fetal bovine serum (FBS) and 1% penicillin-streptomycin. The cells were incubated at 37°C, 5% $CO_2$, and the media was replaced every two days. Cells were passaged at 75-80% confluency using Trypsin-EDTA (0.25%); 1,000,000 HEK 293T cell samples were harvested by detaching the cells with Trypsin-EDTA (0.25%), centrifugation at 700 rcf for 5 minutes, and resuspending in PBS for further experiments. Cell lysate was prepared by mechanical disruption via freeze-thaw cycles.

### 3. Formation of AuNPs assembly

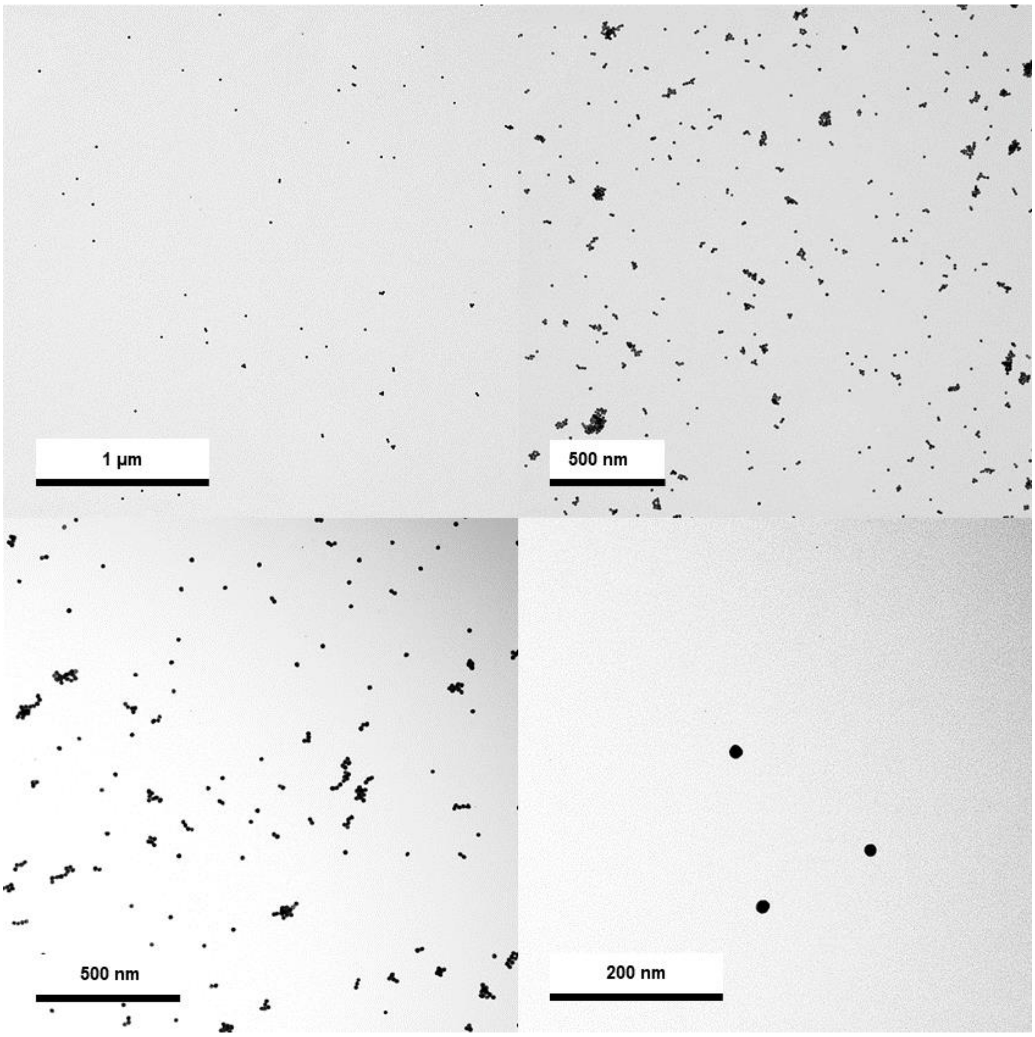

**Figure S1.** TEM images of AuNPs-citrate at different magnifications.

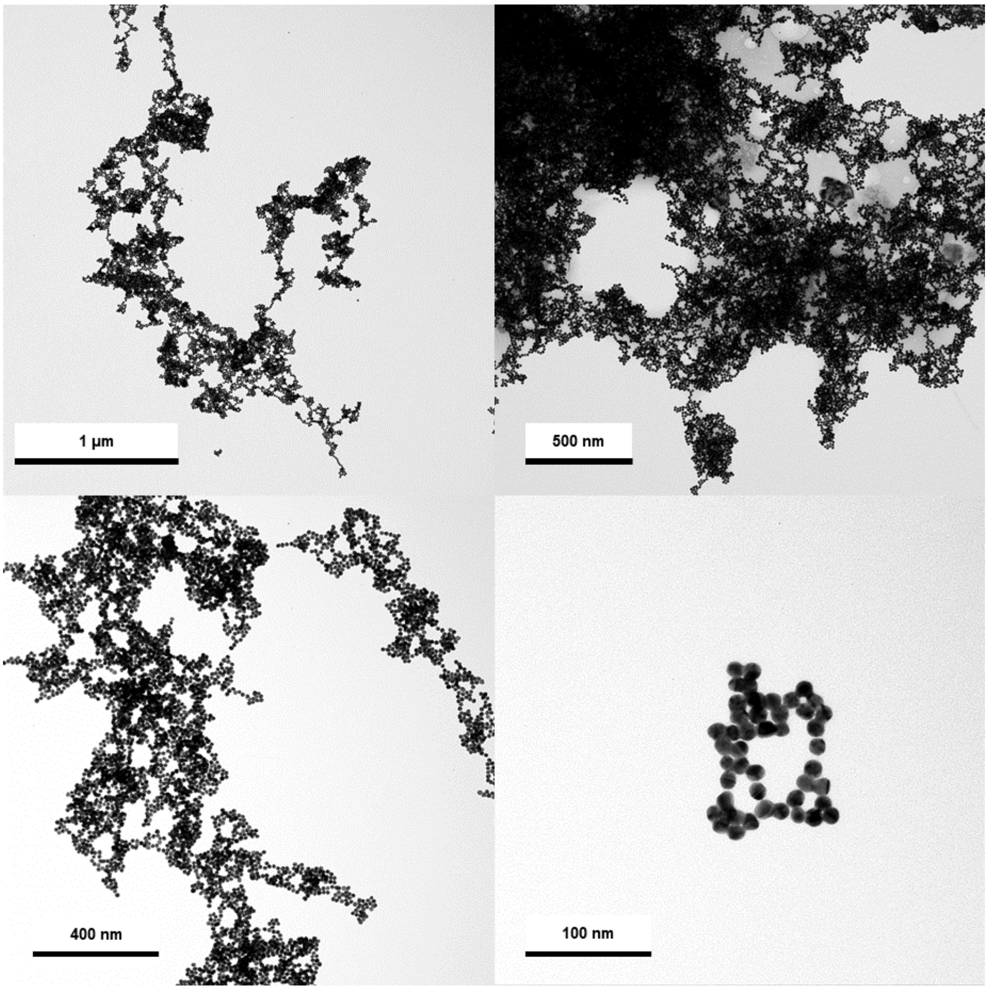

**Figure S2.** TEM images of Arg-Arg-AuNPs at different magnifications.

### 4. Dissociation of the AuNPs assemblies.

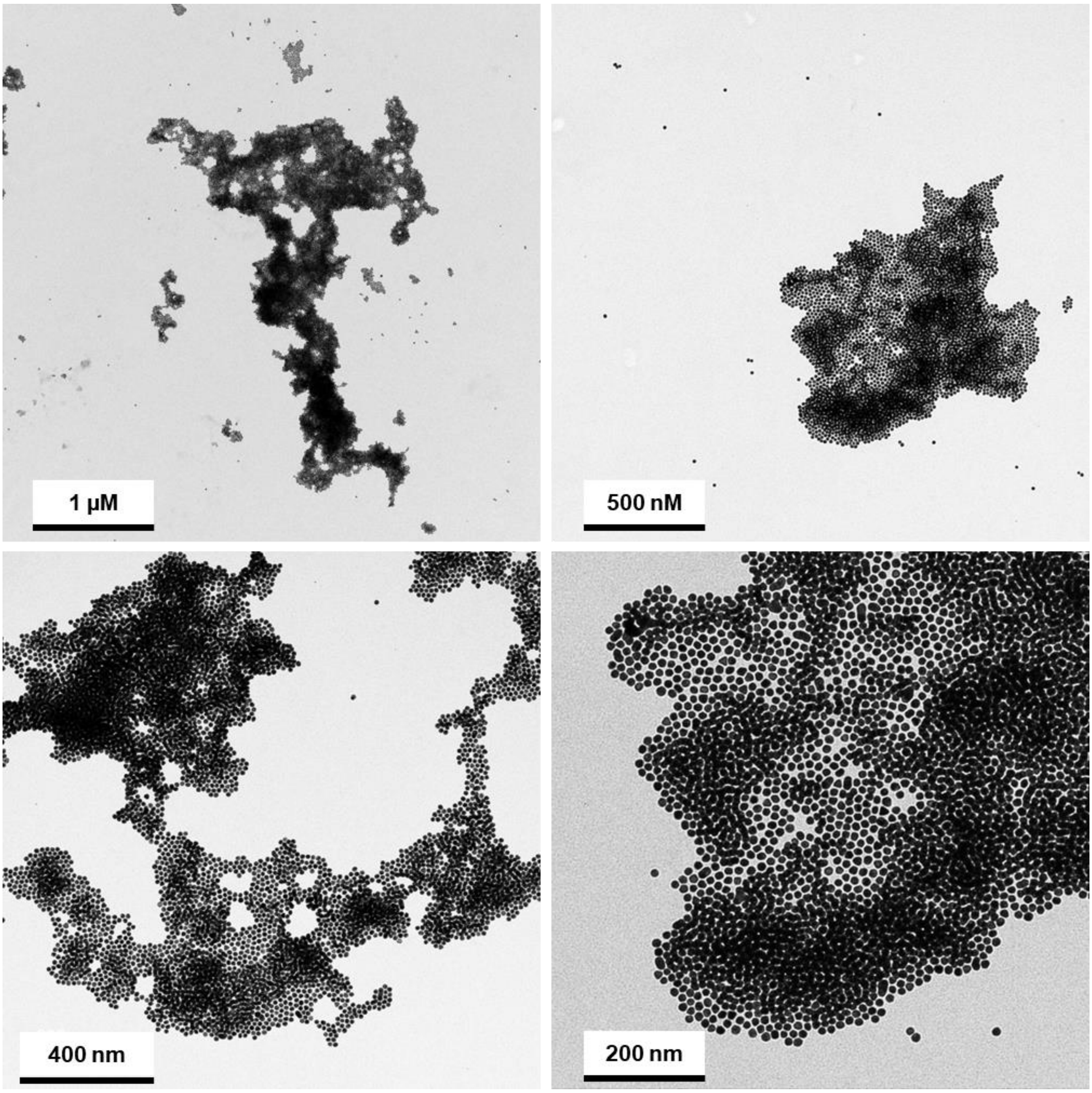

**Figure S3.** TEM images of Arg-Arg-AuNPs after addition of 2 µM of HS-$PEG_6$-$OCH_3$.

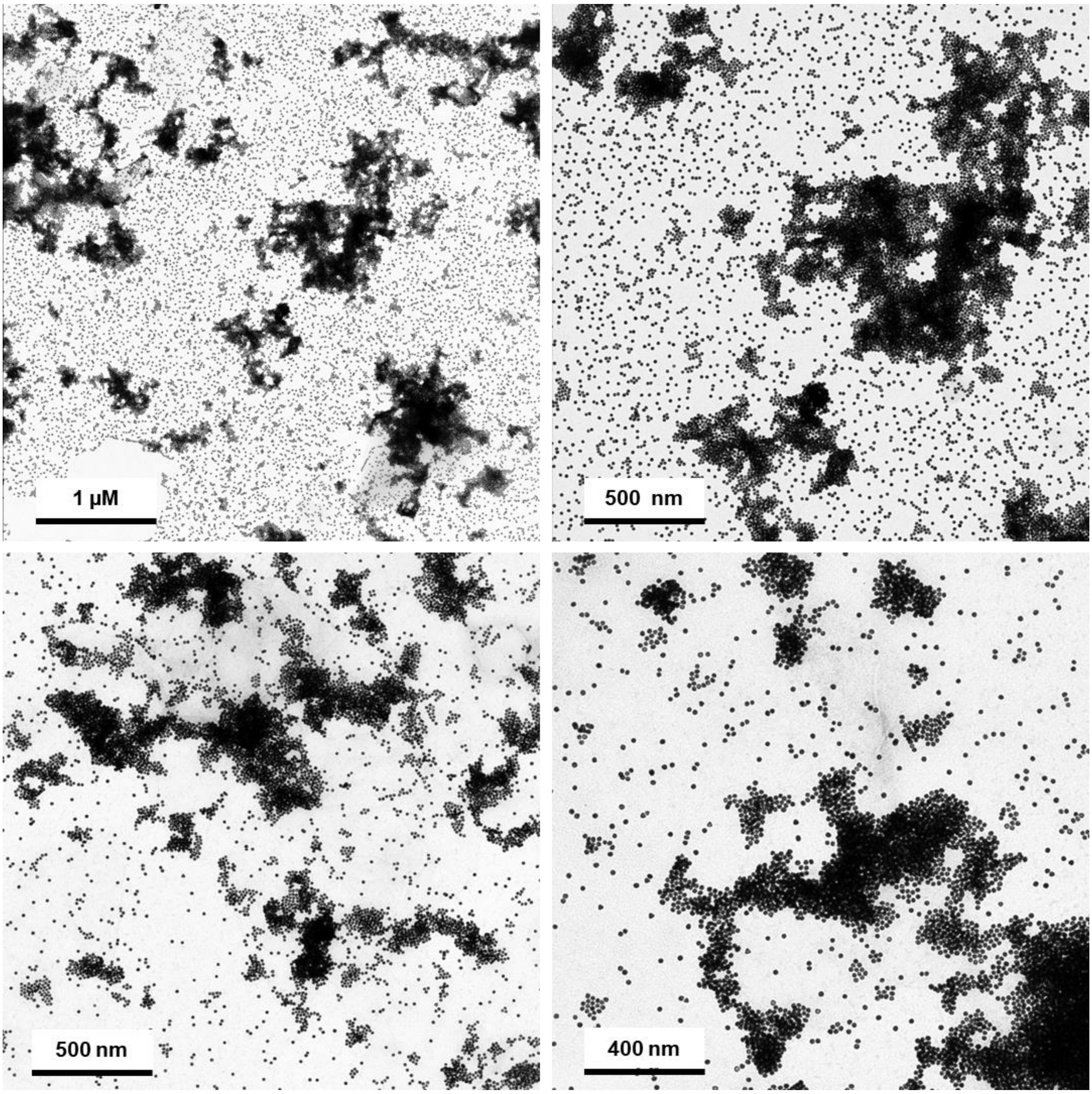

**Figure S4.** TEM images of Arg-Arg-AuNPs after addition of 4 µM of HS-PEG$_6$-OCH$_3$.

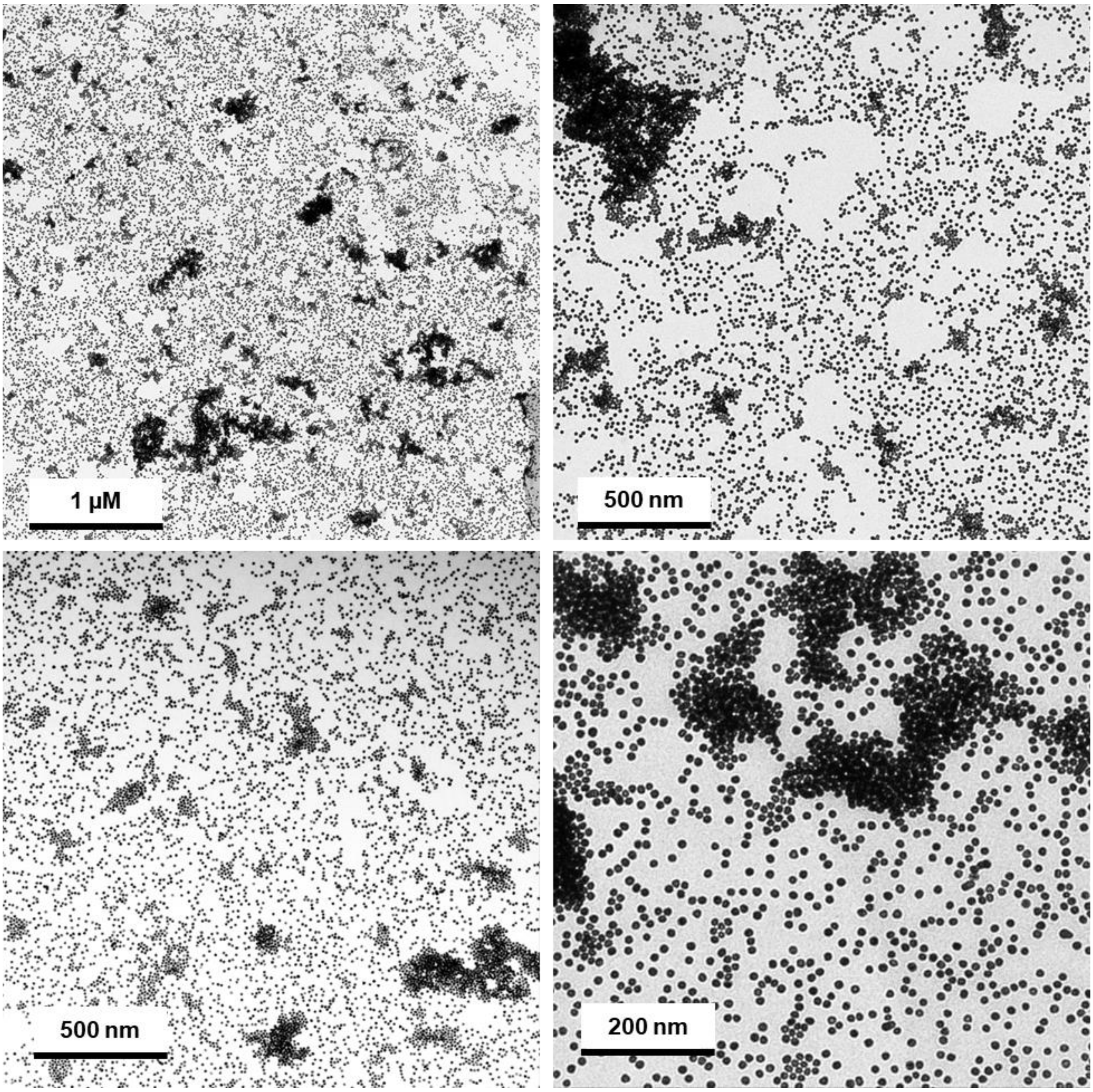

**Figure S5.** TEM images of Arg-Arg-AuNPs after addition of 8 μM of HS-$PEG_6$-$OCH_3$.

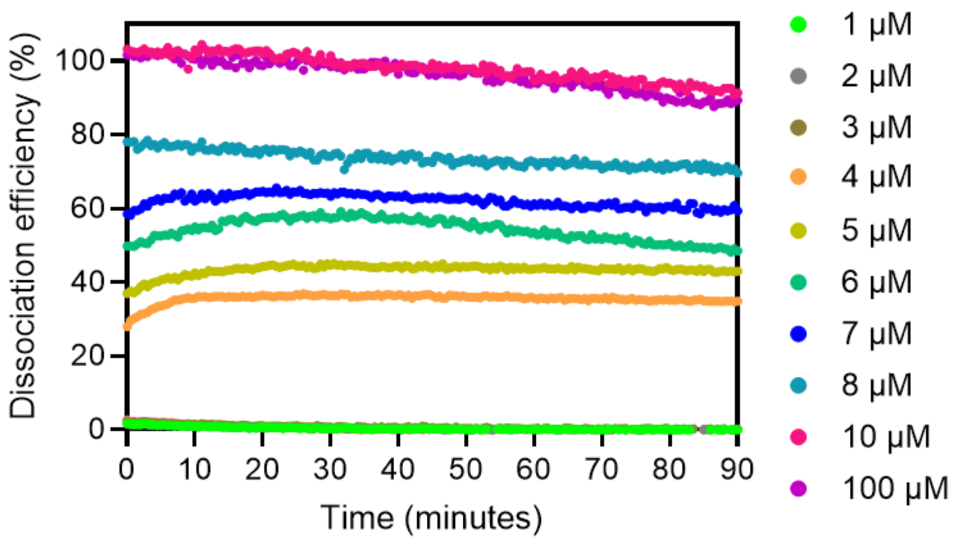

**Figure S6.** Dissociation efficiency over time of HS-PEG$_6$-OCH$_3$ for different concentrations added to Arg-Arg-AuNPs.

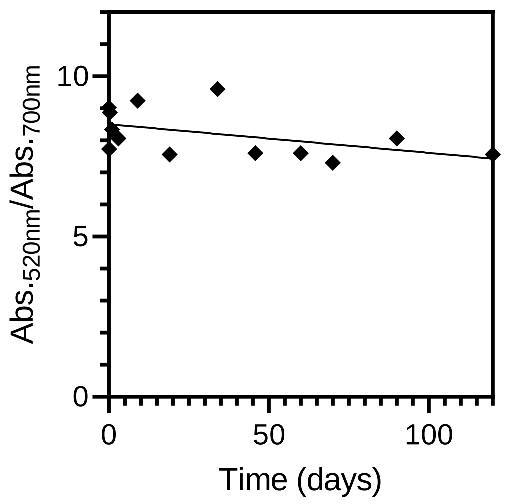

**Figure S7.** Ratio of the absorbances at 520 nm and 700 nm of Arg-Arg-AuNPs dissociated with 10 μM of HS-PEG$_6$-OCH$_3$ as a function of time between the formation of the assembly and the dissociation.

### 5. Dissociation in complex matrices

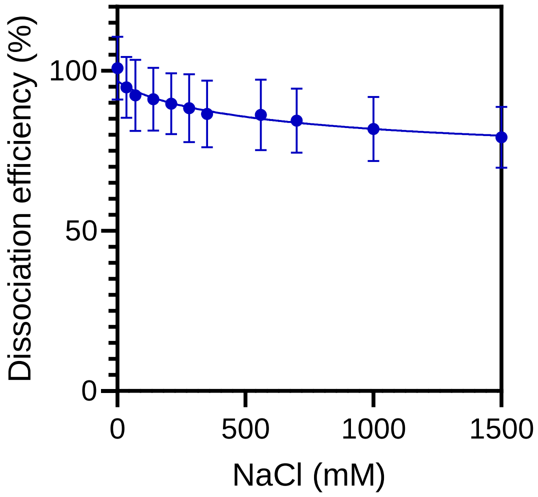

**Figure S8.** Dissociation efficiency of 10 µM of HS-PEG$_6$-OCH$_3$ added to Arg-Arg-AuNPs in increasing NaCl concentrations.

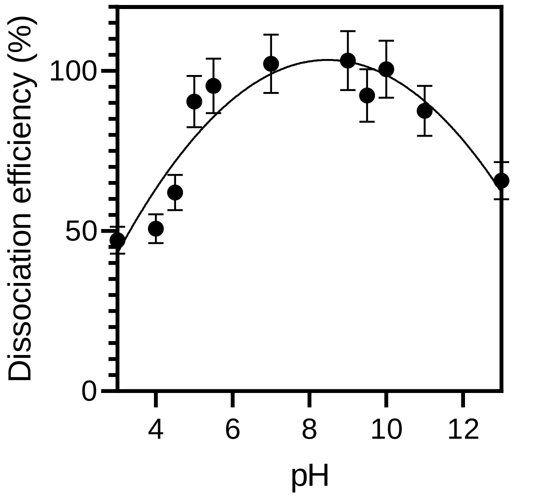

**Figure S9.** Dissociation efficiency of 10 µM of HS-PEG$_6$-OCH$_3$ added to Arg-Arg-AuNPs at different pH values.

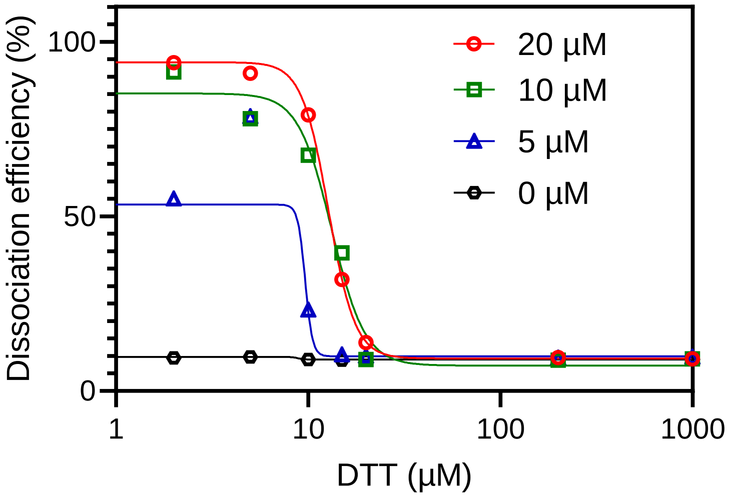

**Figure S10.** Dissociation efficiency of 0, 5, 10 or 20 µM of HS-PEG$_6$-OCH$_3$ added to Arg-Arg-AuNPs in increasing concentration of DTT.

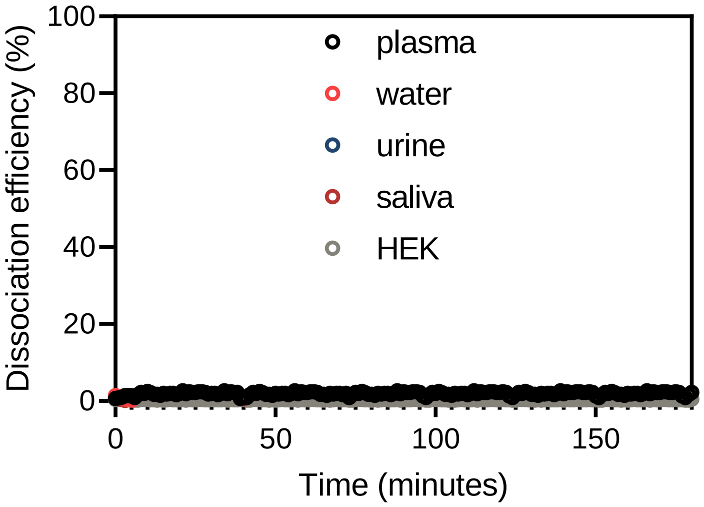

**Figure S11.** Dissociation efficiency for Arg-Arg-AuNPs suspended in different complex matrices over time. HEK = HEK cell lysates coming from suspension of $10^6$ cells/mL in DMEM medium.

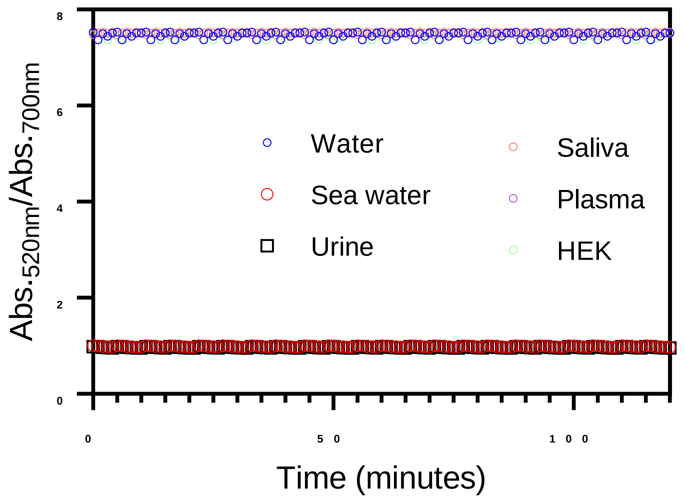

**Figure S12.** Ratio of the absorbances at 520 nm over 700 nm of AuNPs-citrate suspended in various complex matrices. Ratio of AuNPs-citrate dispersed in water =7.8.

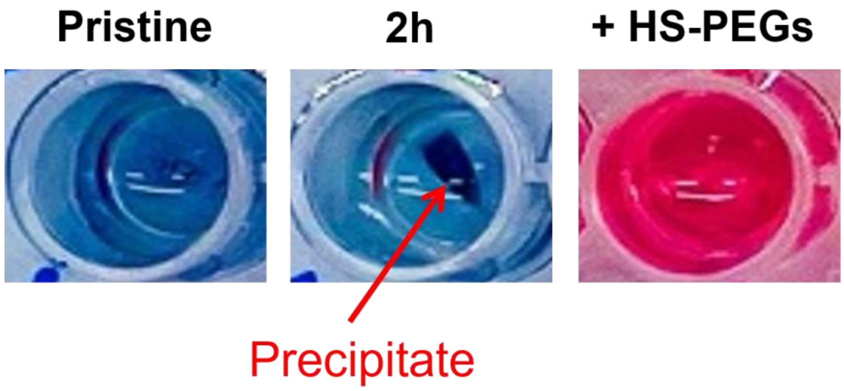

**Figure S13.** Pictures of Arg-Arg-AuNPs suspended in saliva either pristine, after two hours or after two hours and addition of 10 µM of HS-PEG$_6$-OCH$_3$. Red arrows show the formation of the macroscopic precipitates.

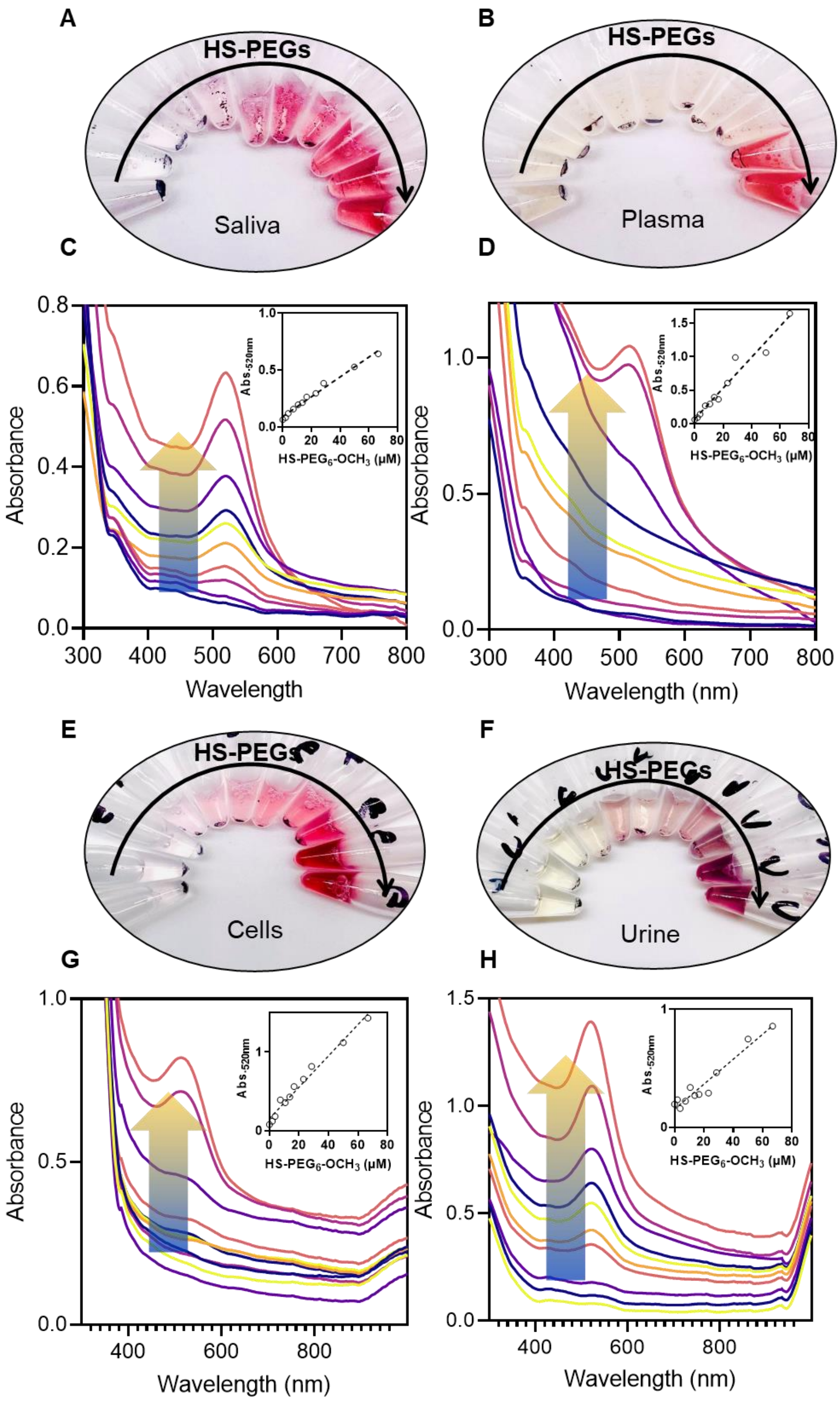

**Figure S14.** Pictures and UV-Vis spectra of dried Arg-Arg-AuNPs film solubilized via the addition of increasing concentrations of HS-PEG$_6$-OCH$_3$ in either **(A)** saliva, **(B)** plasma, **(C)** HEK cell lysates, **(D)** urine. The insets show the absorbance at 520 nm as a function of the HS-PEG$_6$-OCH$_3$ concentration.

### 6. Protease detection

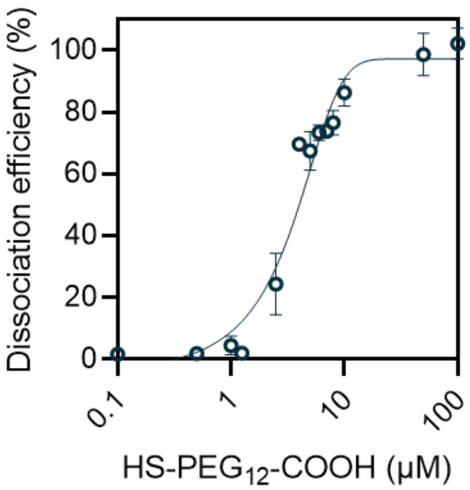

**Figure S15.** Dissociation efficiency of HS-PEG12-COOH (634 Da) added to Arg-Arg-AuNPs.

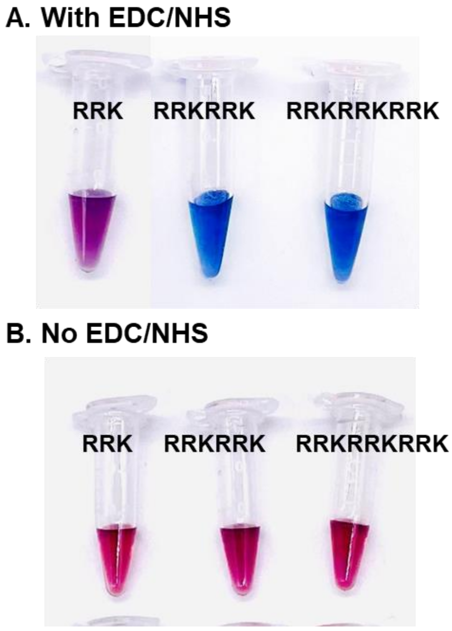

**Figure S16.** Pictures of Arg-Arg-AuNPs 20 minutes after their exposition to 20 µM of HS-PEG12-COOH that was initially mixed with the peptides RRK, RRKRRK or RRKRRKRRK either **(A)** in the presence or **(B)** the absence of EDC/NHS. In the absence of EDC/NHS, there is no conjugation between HS-PEG12-COOH and the peptide and thus, HS-PEG12-COOH can still dissociate the assemblies.

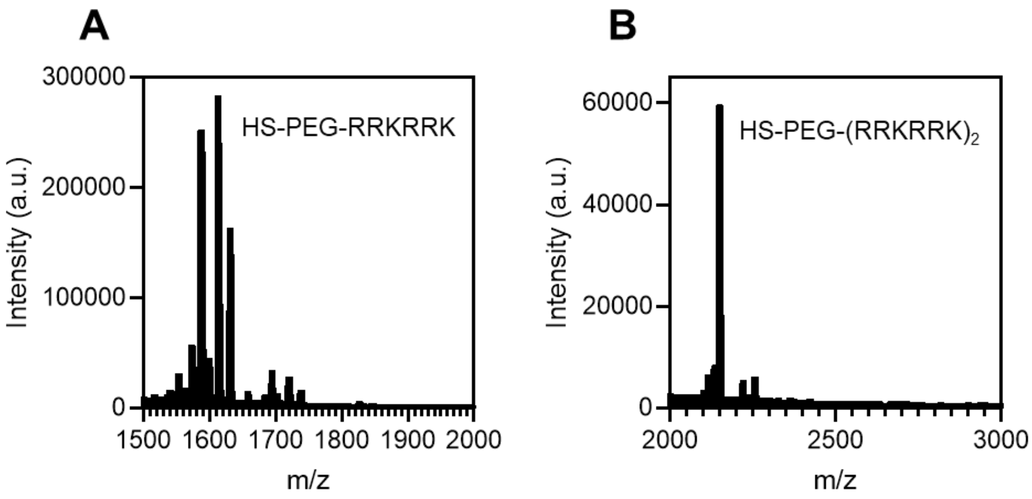

**Figure S17.** Matrix-assisted laser desorption/ionization (MALDI) analysis of HS-PEG-RRKRRK. After the synthesis, the product was purified by HPLC and two compounds could be identified: **(A)** RRKRRK conjugated to one HS-$PEG_{12}$-COOH and **(B)** RRKRRK conjugated to two HS-$PEG_{12}$-COOH units.

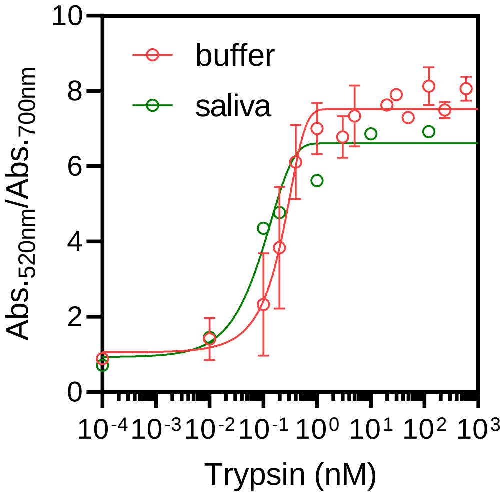

**Figure S18.** Trypsin detection using HS-PEG-RRKRRK either in buffer (PBS 1x, pH 7.4) or in saliva.

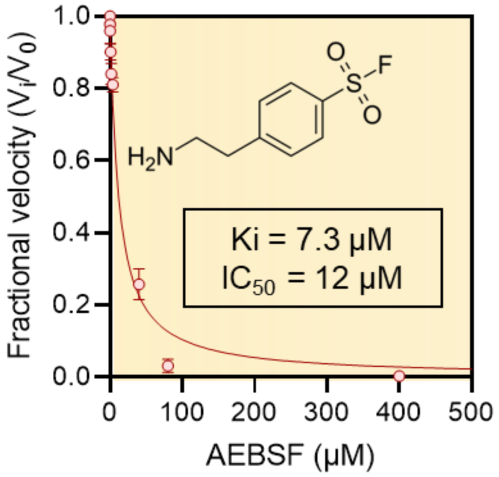

**Figure S19.** A typical inhibition titration curve fitted with the Morrison equation for the competitive inhibitor, AEBSF. Inset shows the chemical structure of AEBSF and the Ki and $IC_{50}$ values. Briefly, 1 μM of trypsin was mixed for 30 minutes at room temperature with increasing concentration of AEBSF. Then, HS-PEG-RRKRRK was added and the resulting solution was incubated at 37.5 °C for 2 hours before addition to Arg-Arg-AuNPs assemblies and the ratio of the absorbance at 520 nm and 700nm was measured.

### 7. Dissociation study with 40 nm AuNPs.

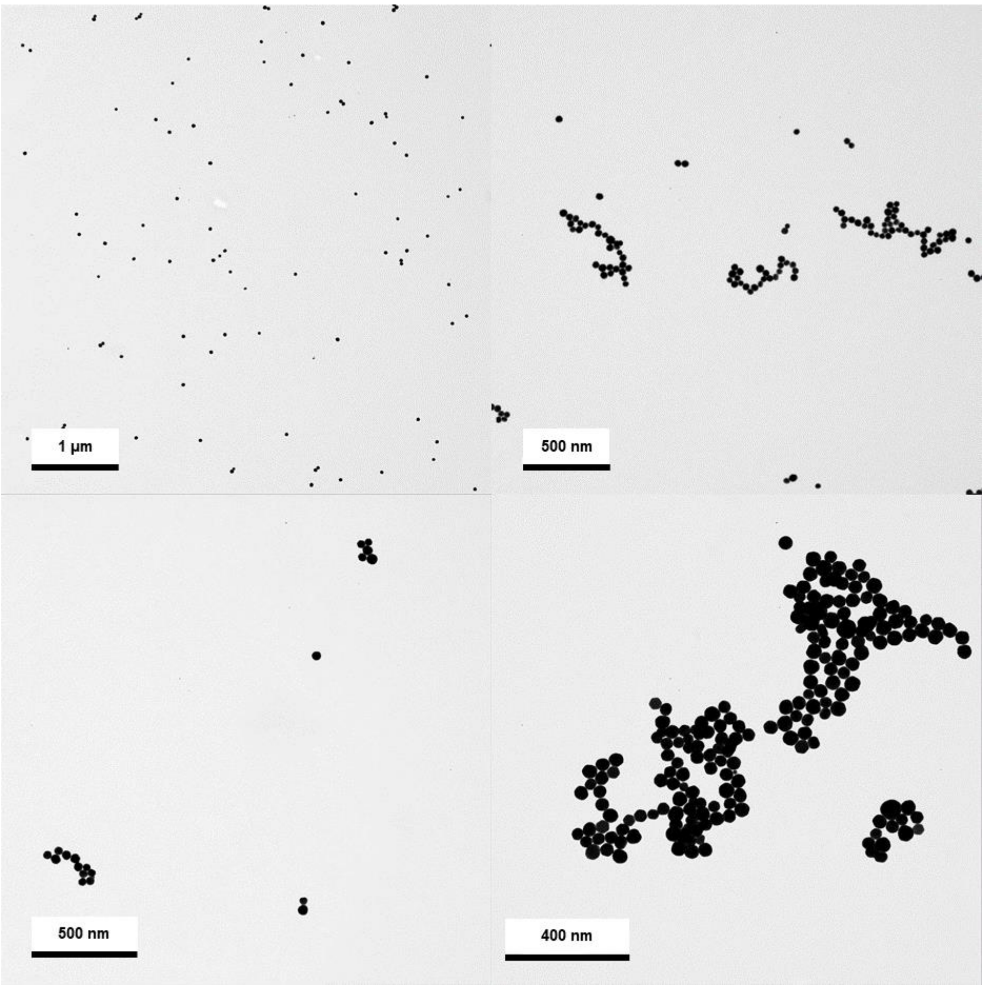

**Figure S20.** TEM images of AuNPs-citrate (40 nm) at different magnifications.

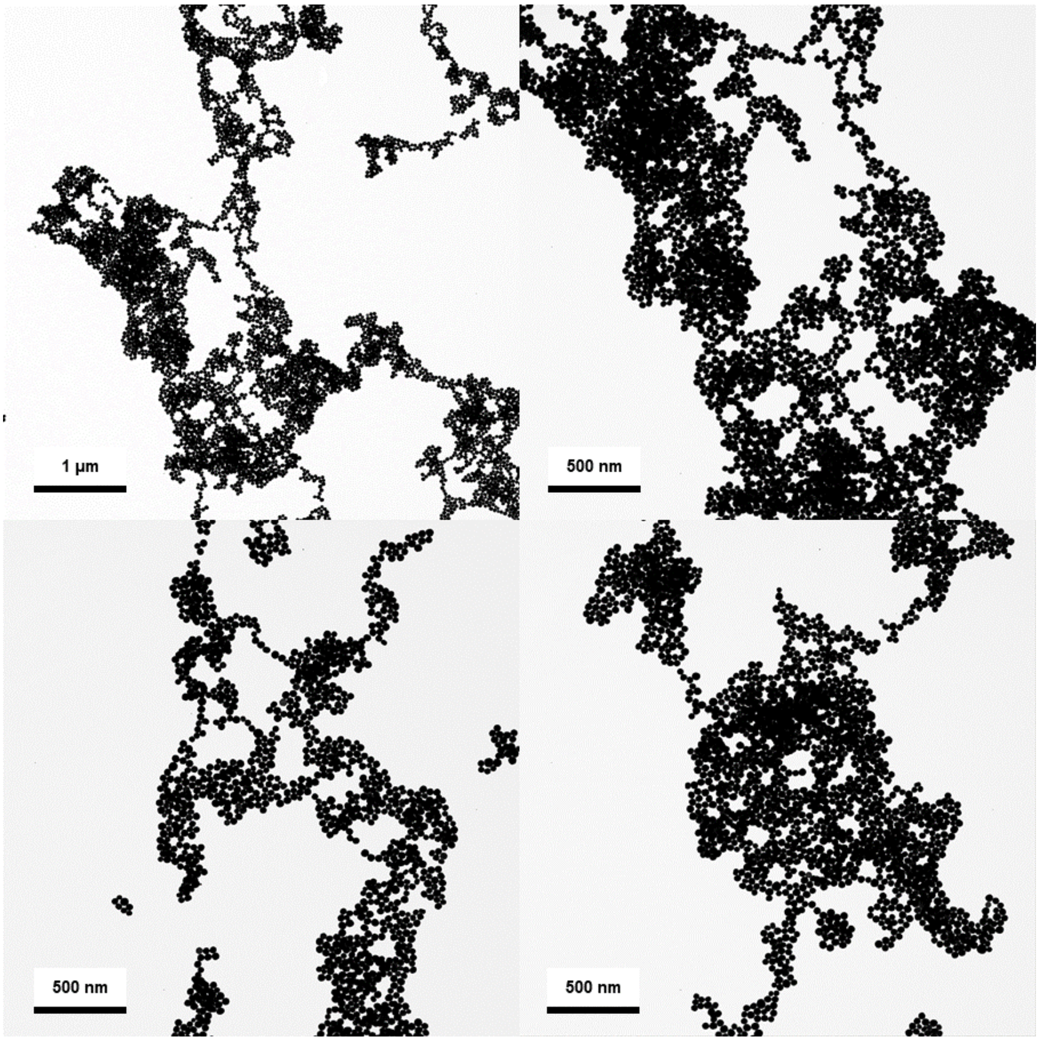

**Figure S21.** TEM images of Arg-Arg-AuNPs (40 nm) at different magnifications.

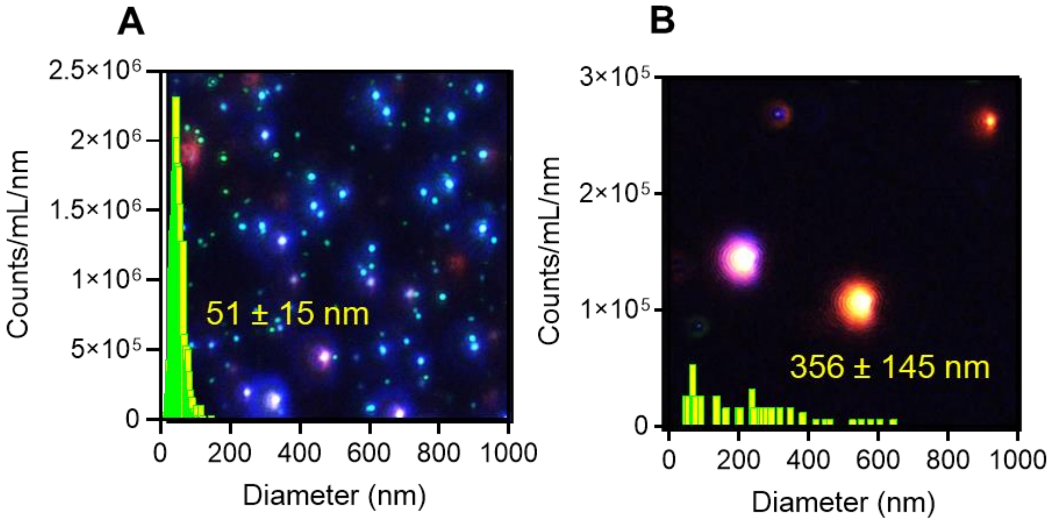

**Figure S22.** Size distribution of AuNPs-citrate (40 nm) **(A)** before and **(B)** 10 minutes after the addition of 2 µM of Arg-Arg obtained via multispectral advanced nanoparticles tracking analysis (MANTA).

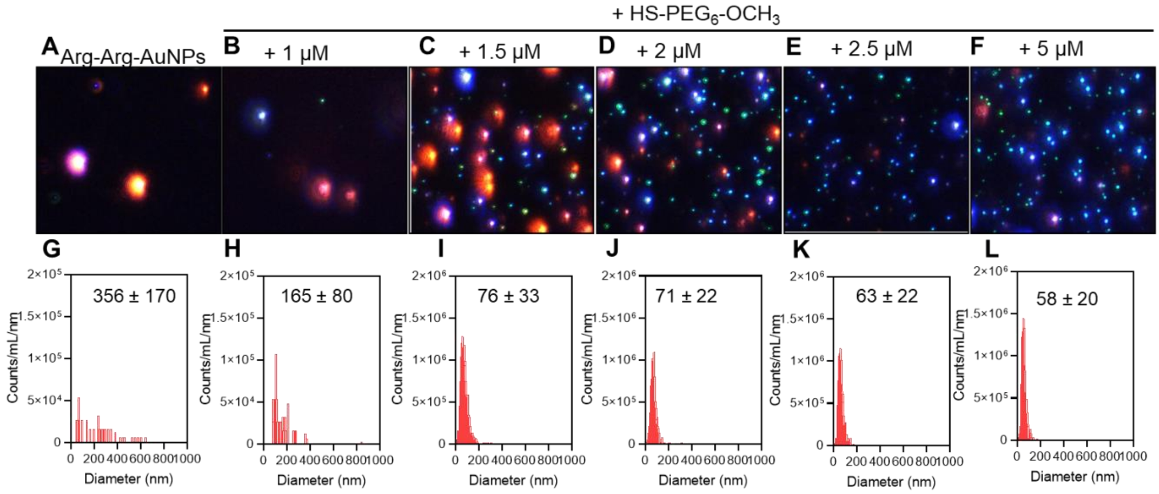

**Figure S23.** MANTA images and size histograms for Arg-Arg-AuNPs (40 nm) (**A** and **G**) 10 minutes after exposure to either 1 µM (**B** and **H**), 1.5 µM, 2 µM, 2.5 µM, or 5 µM (**F** and **L**) of HS-PEG$_6$-OCH$_3$.

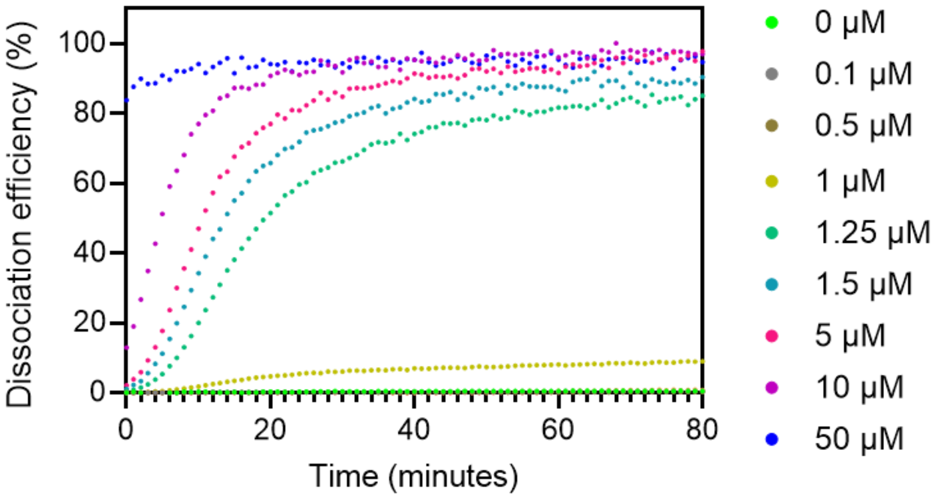

**Figure S24.** Evolution over time of the dissociation efficiency of different concentrations of HS-PEG$_6$-OCH$_3$ added to Arg-Arg-AuNPs (40 nm).

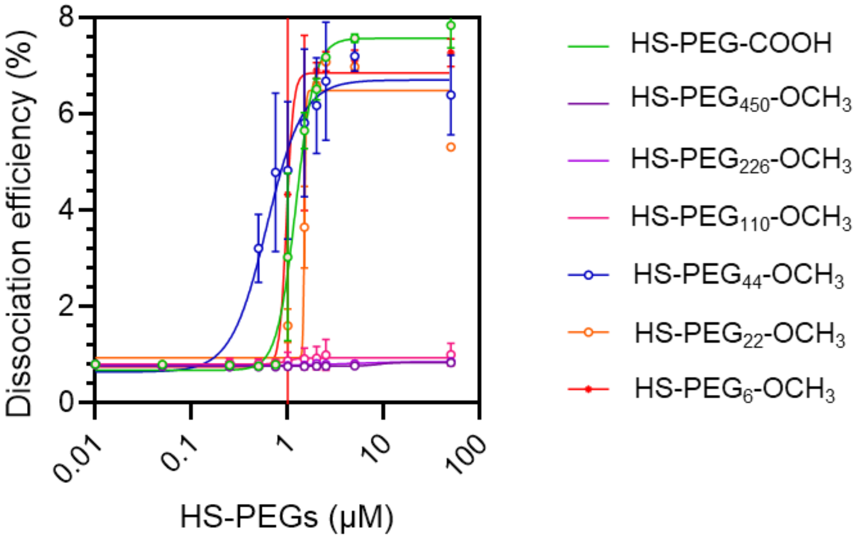

**Figure S25.** Dissociation efficiency of HS-PEGx-OCH$_3$ of different molecular weights added to Arg-Arg-AuNPs (40 nm).

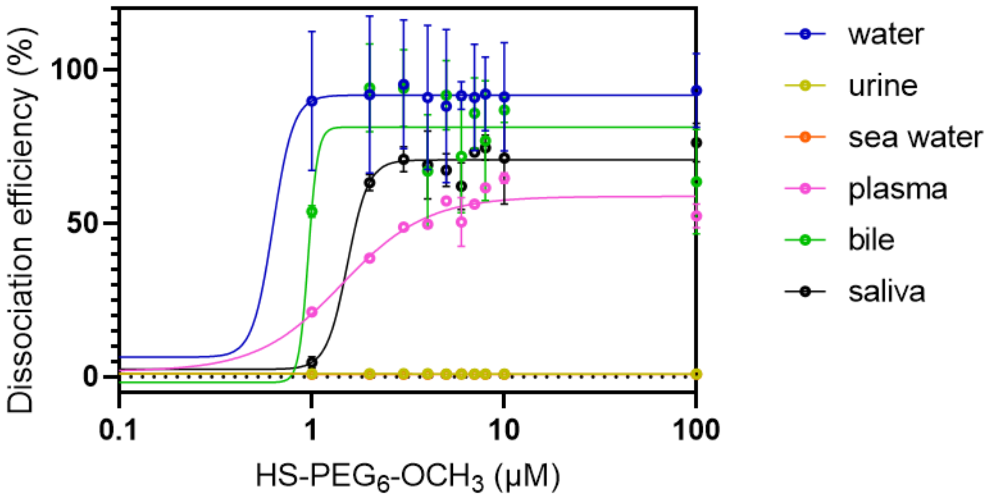

**Figure S26.** Dissociation efficiency (%) of HS-PEG$_6$-OCH$_3$ added to Arg-Arg-AuNPs (40 nm) in different complex matrices.
